# The two-component system ArlRS is essential for wall teichoic acid glycoswitching in *Staphylococcus aureus*

**DOI:** 10.1101/2024.09.10.612275

**Authors:** Marieke M. Kuijk, Emma Tusveld, Esther Lehmann, Rob van Dalen, Iñigo Lasa, Hanne Ingmer, Yvonne Pannekoek, Nina M. van Sorge

## Abstract

*Staphylococcus aureus* is among the leading causes of hospital-acquired infections. Critical to *S. aureus* biology and pathogenesis are the cell wall-anchored glycopolymers wall teichoic acids (WTA). Approximately one-third of *S. aureus* isolates decorates WTA with a mixture of α1,4- and β1,4-*N*-acetylglucosamine (GlcNAc), which requires the dedicated glycosyltransferases TarM and TarS, respectively. Environmental conditions, such as high salt concentrations, affect the abundance and ratio of α1,4- and β1,4-GlcNAc WTA decorations, thereby impacting biological properties such as antibody binding and phage infection. To identify regulatory mechanisms underlying WTA glycoswitching, we screened 1,920 *S. aureus* mutants (Nebraska Transposon Mutant Library) by immunoblotting for differential expression of WTA-linked α1,4- or β1,4-GlcNAc using specific monoclonal antibody Fab fragments. Three two-component systems (TCS), GraRS, ArlRS, and AgrCA, were among the 230 potential hits. Using isogenic TCS mutants, we demonstrated that ArlRS is essential for WTA β1,4-GlcNAc decoration through regulation of *tarM* but not *tarS*. ArlRS regulated *tarM* expression through the transcriptional regulator MgrA, which was responsive to Mg2+, but not Na+. Importantly, ArlRS-mediated regulation of WTA glycosylation affected *S. aureus* interaction with the innate receptor langerin and lysis by β1,4-GlcNAc-dependent phages. Since WTA represents a promising target for future immune-based treatments and vaccines, our findings provide important insight to align targeting strategies with the *S. aureus* WTA glycosylation patterns during infection.

**Importance:** *Staphylococcus aureus* is a common colonizer of mucosal surfaces, but is also a frequent cause of severe infections in humans. Development of antibiotic resistance complicates treatment of *S. aureus* infections, increasing the need for antibiotic alternatives such as vaccines and therapies with bacterial viruses also known as phages. Wall teichoic acids (WTA) are abundantly-expressed glycosylated structures of the *S. aureus* cell wall that have gained attention as a promising target for new treatments. Importantly, WTA glycosylation patterns show variation depending on environmental conditions, thereby impacting phage binding and interaction with host factors, such as antibodies and innate pattern-recognition receptors. Here, we show that the two-component system ArlRS through its effector MgrA is involved in regulation of WTA glycosylation by responding to environmental changes in Mg^2+^ concentration. These findings may support the design of new treatment strategies that target WTA glycosylation patterns of *S. aureus* during infection.

## Introduction

Antimicrobial resistance is a global health crisis, which is estimated to cause an increasing number of deaths worldwide in the upcoming years [1]. Infections caused by the Gram-positive pathogen *Staphylococcus aureus* are associated with high morbidity and mortality rates, and methicillin-resistant *S. aureus* (MRSA) specifically accounted for more than 100,000 deaths globally in 2019 [1]. Consequently, *S. aureus* is one of six pathogens that significantly contributes to the number of multidrug resistant deaths and is classified by the WHO as a high priority pathogen [2]. Novel treatments to combat these infections are therefore urgently needed.

Wall teichoic acids (WTA) are critical glycopolymers for *S. aureus* cell wall architecture, physiology and host interaction. WTA molecules are covalently anchored to peptidoglycan and are composed of polyribitol-phosphate decorated with *N*-acetylglucosamine (GlcNAc) and D-alanine. WTA glycosylation plays an important role in a wide range of processes, including β-lactam resistance [3], phage infectivity [4, 5], and host interactions such as nasal colonization [6, 7], detection by WTA-specific antibodies [8–10] as well as the langerin receptor on skin Langerhans cells [11]. The GlcNAc residues can be linked in several distinct orientations, resulting in structural heterogeneity of WTA. Nearly all *S. aureus* isolates glycosylate WTA with β1,4-GlcNAc through the activity of the specific house-keeping glycosyltransferase TarS [3, 12]. However, approximately one-third of *S. aureus* isolates can co-decorate WTA with α1,4-GlcNAc through the activity of the accessory glycosyltransferase TarM [12, 13]. The *tarS* gene is part of the core genome and is co-localized with several other WTA biosynthesis genes, whereas *tarM* is located elsewhere in the genome [3, 14]. Importantly, several of the WTA-mediated processes, e.g. β-lactam resistance, langerin binding, and infection by specific phages, are dependent on WTA β1,4-GlcNAc and are blocked by co-decoration with α1,4-GlcNAc [15–17]. The β1,4-GlcNAc moiety is also a dominant target for human antibodies and is therefore an interesting target for immune-mediated treatments and vaccines [8, 10, 18]. Clearly, these findings illustrate how changes in WTA glycosylation affect and shape host-pathogen interactions.

*S. aureus* can rapidly adapt its surface properties to different environmental conditions encountered during its commensal and pathogenic lifestyles, which is a major challenge for treating infections as well as vaccine development [19, 20]. One of these adaptations concerns the shift in the abundance of α1,4-GlcNAc and β1,4-GlcNAc WTA glycosylation. For example, bacteria grown *in vitro* under rich growth conditions primarily decorate WTA with α1,4-GlcNAc, whereas *S. aureus* isolated from organs after murine infection or after culture in high salt conditions expresses predominantly β1,4-GlcNAc WTA [8, 21]. Culture density also affects α1,4-/β1,4-GlcNAc WTA glycosylation ratios through quorum sensing, as auto-inducing peptides have been observed to reduce *tarM* expression in stationary phase cultures, resulting in increased lysis by b-GlcNAc-dependent phages [22]. Although these examples illustrate the ability and relevance of *S. aureus* WTA glycoswitching, the underlying molecular mechanisms and regulatory pathways are currently unknown.

*S. aureus* is able to quickly adapt its gene expression profile [23, 24] through a network of transcriptional regulators (SarA, Rot, MgrA, among others), alternative sigma factors (SigB and SigH) [23] and two-component systems (TCS) [25]. *S. aureus* encodes 16 TCS, of which only WalRK is essential [25, 26]. TCS respond to specific external triggers by a sensing histidine kinase, leading to transcriptional regulation through the phosphorylated response regulator [27]. Intriguingly, no major growth defects were observed in an *S. aureus* mutant lacking all 15 non-essential TCS under standard laboratory conditions [25]. However, many TCS, such as AgrCA, SaeRS, SrrAB and ArlRS, are important for bacterial virulence or survival in certain *in vivo* conditions [23, 27]. With regard to WTA glycosylation, only AgrCA has been experimentally confirmed to affect WTA glycosylation profiles through the transcriptional regulation of *tarM* [22]. GraRS has also been implicated in *tarM* regulation [28], but WTA glycosylation levels were not analyzed to confirm a functional effect. Finally, the TCS ArlRS has been linked to WTA, but as a repressor of the *dlt* operon encoding the machinery for WTA D-alanylation [29]. In general, ArlRS is important in regulating host interaction such as adhesion and damage to endothelial cells [30, 31], clumping with fibrinogen [31–33], and expression of several immune evasion factors that block neutrophil killing [34]. Although the exact triggers for activation of ArlRS in *S. aureus* have not been identified, ArlRS has been implicated in homeostasis of the essential divalent cations Mn^2+^ and Mg^2+^ [29, 35, 36].

Despite the implication of TCS, the regulatory mechanisms that control WTA glycoswitching in response to varying environmental conditions remain largely unknown. This study aimed to identify key regulators and unravel the molecular mechanisms involved in differential expression of α1,4- and β1,4-GlcNAc phenotypes. We demonstrate that ArlRS is essential for β1,4-GlcNAc decoration of WTA, which impacts phage infection and langerin interaction. In addition, ArlRS was required for the Mg^2+^-induced glycoswitch from α1,4- to β1,4-GlcNAc WTA glycosylation via the transcription factor MgrA. Since WTA represents a promising target for future immune-based treatments and vaccines, our findings are important to align targeting strategies with the WTA glycosylation patterns of *S. aureus* during infection.

## Results

### Environmental conditions affect *tarM*, but not *tarS*, transcription levels

*S. aureus* strains containing *tarM* and *tarS* can co-decorate WTA with α1,4- and β1,4-GlcNAc, respectively. To understand the molecular mechanisms underlying the regulation of the α1,4- and β1,4-GlcNAc ratio [21], we investigated how *tarM* and *tarS* expression levels were affected in different environments. We therefore re-analyzed a previously published transcriptomics dataset where *S. aureus* strain HG001 was exposed to 44 different conditions ranging from *in vitro* lab culture to *in vivo*-mimicking host environments [24]. We extracted transcription data for the *tarM* and *tarS* genes [37] and compared the changes in expression levels of the two genes across these conditions. *tarM* expression fluctuated substantially, which was significantly different from the stable *tarS* expression in these 44 experimental conditions (Figure 1). These results suggest that shifts in the WTA glycosylation profile are predominantly regulated through *tarM* rather than *tarS*.

**Figure 1.**
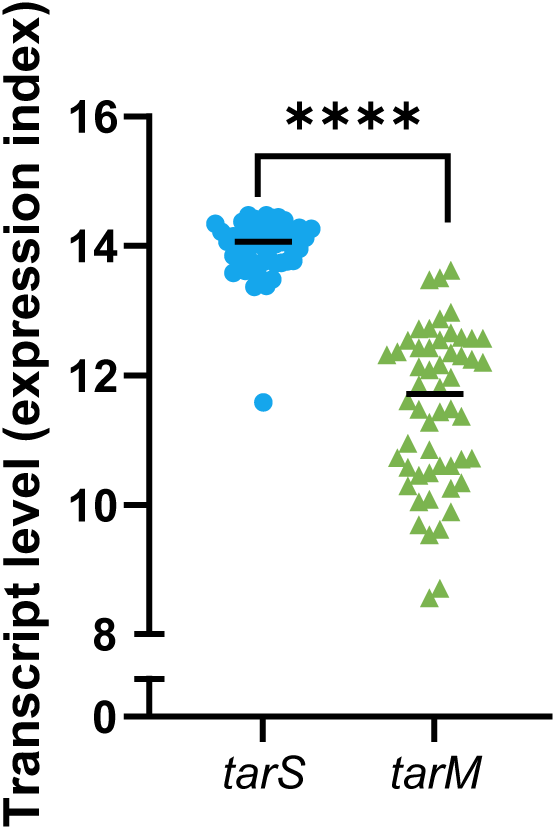
*tarM* expression is dynamically regulated. Transcript levels of *tarS* (SAOUHSC_00228) and *tarM* (SAOUHSC_00973) of *S. aureus* HG001 in 44 different *in vitro* and *in vivo*-mimicking conditions. Data are extracted from Mäder *et al*. [24, 37]. The variance of the two genes was statistically tested using an F test. \*\*\*\**p* < 0.0001.

### Multiple two-component systems are identified to be involved in WTA glycoswitching

Given the dynamic regulation of *tarM*, we next aimed to identify the genes that may be involved in this process. To this end, we screened the Nebraska Transposon Mutant Library (NTML), containing 1,920 arrayed mutants, for differential α1,4- and β1,4-GlcNAc expression [38]. Importantly, the NTML parental strain JE2 harbors both *tarM* and *tarS* genes. Individual transposon mutants were screened by immunoblotting (Figure 2A) in two parallel screens using Fab fragments that specifically detect α1,4- and β1,4-GlcNAc WTA [39].

**Figure 2.**
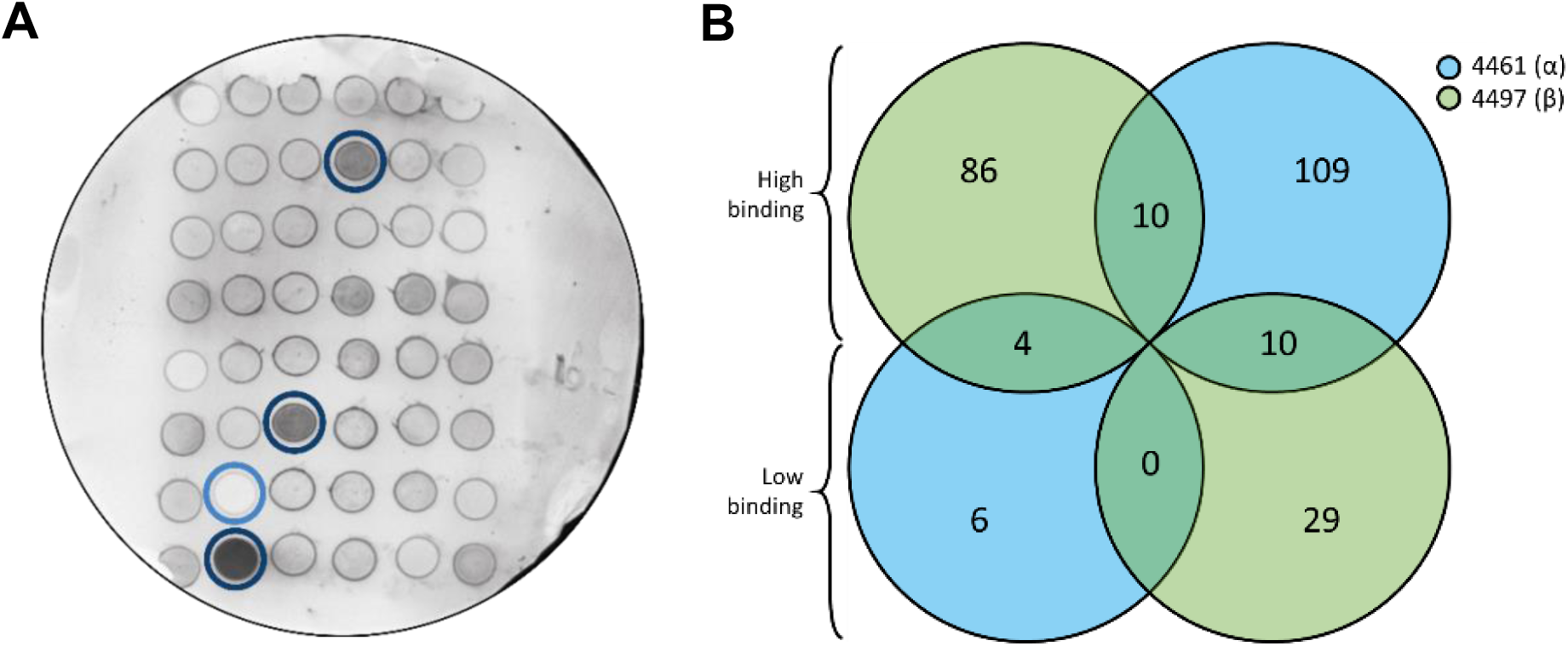
Representative immunoblot and Venn diagram summarizing results from the NTML screen to identify mutants with altered WTA glycoprofile. (A) Representative image of an immunoblot containing 48 NTML mutants screened for binding of 4461 (α1,4-GlcNAc). Three transposon mutants show comparatively high binding (encircled in dark blue) and one transposon mutant show low binding (encircled in light blue). (B) Venn diagram with number of NTML mutants identified with either high or low levels of α1,4-(in blue) and β1,4-glycosylation (in green). The number of overlapping mutants is also depicted.

We identified 230 genes potentially involved in WTA glycosylation (Table S3). Importantly, the transposon mutants *tarM* (SAUSA300_0939) and *tarS* (SAUSA300_0252) were identified as non-binding mutants in their respective α1,4- and β1,4-GlcNAc screens, thereby acting as positive controls of this unbiased screening method. Among the 230 potential genes, 115 were identified in the α1,4-GlcNAc and 115 in the β1,4-GlcNAc screen (Figure 2B, Table S3). A higher number of transposon mutants exhibited upregulated WTA glycosylation compared to the number of mutants in which WTA glycosylation was downregulated (195 compared to 35; Figure 2B). Ten mutants showed high binding to both Fabs, whereas 14 genes had a differential effect on α1,4- and β1,4-GlcNAcylation (Figure 2B). Of interest, we identified six genes of three different TCS (GraRS, ArlRS and AgrCA) in the 4461 (α1,4-GlcNAc) screen (Table 1). In addition, nine transcriptional regulators were identified across both screens (Table 1).

**Table 1:** Identified transposon mutants in two-component systems and regulatory proteins.

| Accession No. | NTML ID | Gene | Description | Fab binding |  |
| --- | --- | --- | --- | --- | --- |
|  |  |  |  | 4461 (α) | 4497 (β) |
| Two-component systems |  |  |  |  |  |
| SAUSA300_0646 | NE1756 | *graS* | Sensor protein kinase GraS | High | - |
| SAUSA300_1307 | NE1183 | *arlS* | sensor histidine kinase protein | High | - |
| SAUSA300_1308 | NE1684 | *arlR* | DNA-binding response regulator | High | - |
| SAUSA300_1989 | NE95 | *agrB* | accessory gene regulator protein B | High | - |
| SAUSA300_1991 | NE873 | *agrC* | accessory gene regulator protein C | High | - |
| SAUSA300_1992 | NE1532 | *agrA* | accessory gene regulator protein A | High | - |
| Regulators |  |  |  |  |  |
| SAUSA300_0114 | NE165 | *sarS* | staphylococcal accessory regulator | Low | High |
| SAUSA300_0605 | NE1193 | *sarA* | accessory regulator A | High | - |
| SAUSA300_0647 | NE645 | *vraF* | ABC transporter, ATP-binding protein | High | - |
| SAUSA300_1708 | NE386 | *rot* | staphylococcal accessory regulator Rot | Low | High |
| SAUSA300_2022 | NE1109 | *rpoF* | RNA polymerase sigma factor SigB | High | High |
| SAUSA300_2025 | NE1607 | *rsbU* | sigma-B regulation protein | High | High |
| SAUSA300_2326 | NE1304 | - | transcription regulatory protein | Low | - |
| Other |  |  |  |  |  |
| SAUSA300_0023 | NE1865 | *yycI* | YycI protein (regulation WalRK) | High | - |
| SAUSA300_2075 | NE149 | *rho* | transcription termination factor<br>(regulation SaeRS) | - | High |

Glycosylation patterns of NTML mutants were assessed with either high, low or average (-) levels of Fab binding of 4461 ( α1,4-GlcNAc) or 4497 ( β1,4-GlcNAc).

### The two-component system ArlRS is essential for WTA β-GlcNAc expression

Based on identification of three different TCS, we next aimed to confirm the role of these TCS in regulating WTA glycosylation. We first analyzed a deletion mutant lacking all 15 non-essential TCS (ΔXV) [25], and observed that this mutant completely lacked WTA β1,4-GlcNAc glycosylation (Figure 3A). Further analysis of three isogenic TCS mutants, Δ*arlRS,* Δ*agrCA, and* Δ*graRS*, showed that the WTA glycosylation phenotype of ΔXV mutant was mimicked by the Δ*arlRS* mutant with regard to β1,4-glycosylation, although this mutant additionally showed significantly increased α1,4-GlcNAc levels (Figure 3A). In contrast to *arlRS*, deletion of *graRS* and *agrCA* did not significantly alter α1,4- or β1,4-GlcNAc WTA glycosylation levels (Figure 3A). The β1,4-GlcNAc-deficient phenotype of ΔXV and Δ*arlRS* was completely rescued through *arlRS* plasmid complementation (Figure 3B), demonstrating that *arlRS* is essential for the expression of WTA β1,4-GlcNAc expression in our assay conditions.

**Figure 3.**
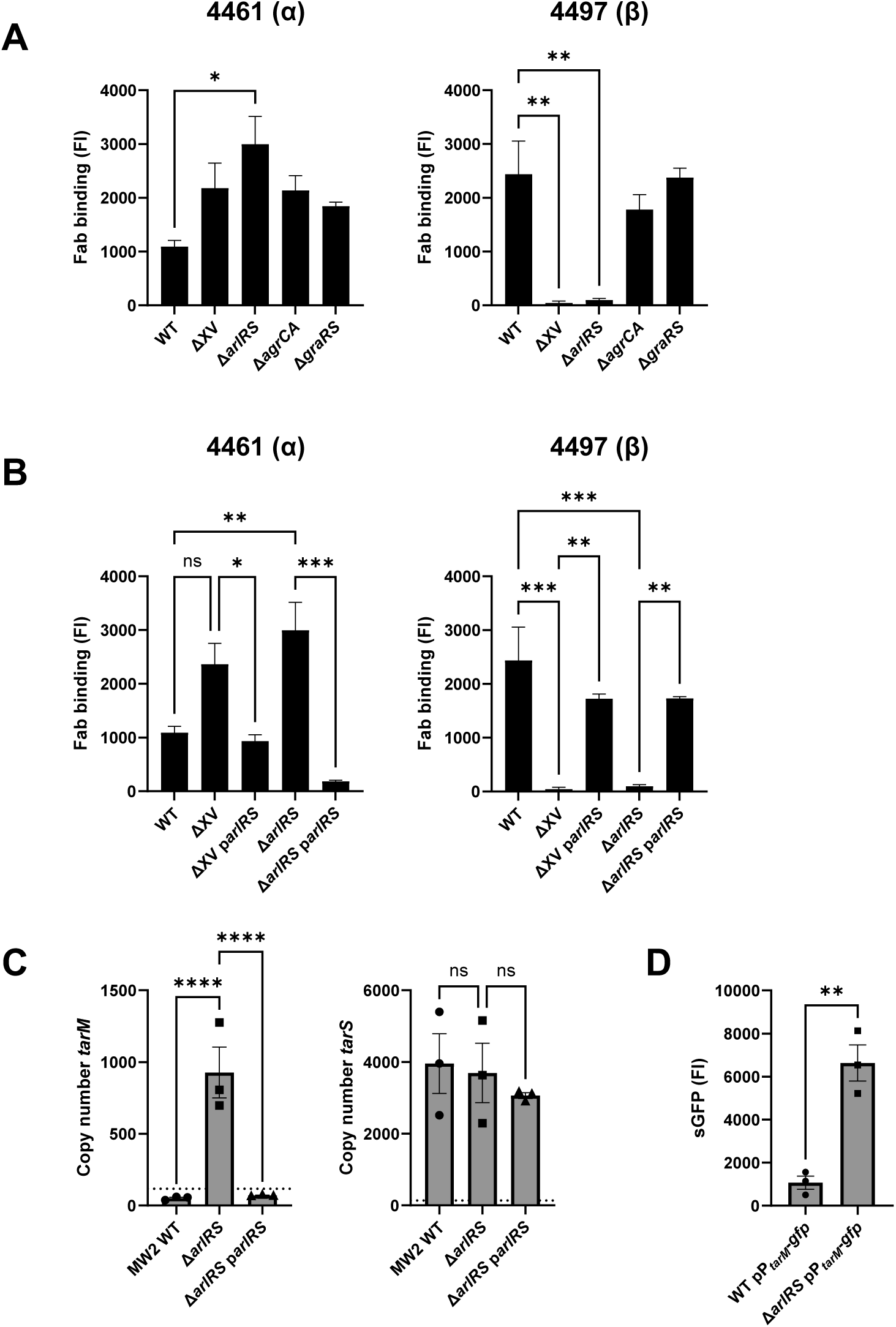
The two-component system ArlRS is essential for WTA β1,4-GlcNAc glycosylation. Levels of α1,4- and β1,4-GlcNAc, quantified by binding of Fab fragments 4461 (α) and 4497 (β) to (A) WT, ΔXV and three single TCS mutants Δ*arlRS*, Δ*agrCA* and Δ*graRS*, and to (B) WT, ΔXV and Δ*arlRS* and their *arlRS* plasmid-complemented strains. All mutants are in the *S. aureus* MW2 background. Fab binding was analyzed by flow cytometry. (C) mRNA copy number of *tarM* and *tarS* as measured with qPCR in WT, Δ*arlRS* and complemented bacteria. Symbols below the dotted line represent extrapolated values. (D) sGFP fluorescence as a measure for promoter activity of *tarM* in WT pP*_tarM_*-*gfp* or Δ*arlRS* pP*_tarM_*-*gfp* bacteria as analyzed by flow cytometry and represented as fluorescence intensity (FI). Data are shown as three biological replicates ± SEM. Statistical significance was determined using one-way ANOVA with Bonferroni statistical hypothesis testing to correct for multiple comparisons (A, B, C) and Student’s t test (D). For panel (A), the mean of each column was compared to WT marking only significant comparisons. \**p* < 0.05, \*\**p* < 0.01, \*\*\**p* < 0.001, \*\*\*\**p* < 0.0001, ns=not significant.

To investigate whether ArlRS regulated expression of *tarM* or *tarS*, we measured the absolute copy numbers of *tarM* and *tarS* by qPCR. The Δ*arlRS* mutant showed a substantial increase in *tarM* copy numbers compared to wild type (WT), which was reversed by complementing the mutant with plasmid-expressed *arlRS.* In contrast, no changes were observed between the *tarS* copy numbers of WT, Δ*arlRS*, and the complemented mutant (Figure 3C). To assess whether increased *tarM* copy numbers correlated with increased promoter activity, we used an sGFP-reporter system in which the *tarM* promoter region was fused to *gfp* in plasmid pCM29 [40]. This plasmid was transformed into WT and Δ*arlRS* to create WT pP*_tarM_*-*gfp* and Δ*arlRS* pP*_tarM_*-*gfp*. Δ*arlRS* showed a significantly increased *tarM* promoter activity compared to WT (Figure 3D), indicating that ArlRS acts as a repressor of *tarM* expression, which promotes WTA β1,4-GlcNAc decoration.

### High Mg^2+^ concentrations drive WTA glycoswitching through ArlRS

High salt concentrations are known to induce a shift in *S. aureus* WTA glycosylation patterns from predominantly α1,4-GlcNAc to β1,4-GlcNAc decoration [21]. The high salt medium previously used contained both Mg^2+^ (15 g/L MgCl_2_ = 158 mM) and Na^+^ (41 g/L NaCl = 702 mM) to induce a stress response. Intriguingly, ArlRS activity was linked to the presence of the divalent cation Mg^2+^ [29]. To investigate whether the salt-induced WTA glycoswitch depended on Mg^2+^ or Na^+^, WT and Δ*arlRS* were grown in TSB and in TSB supplemented with either 200 mM Mg^2+^ (TSB+Mg^2+^) or 200 mM Na^+^ (TSB+Na^+^). WTA glycosylation was assessed by binding of Fabs 4461 (α1,4-GlcNAc) and 4497 (β1,4-GlcNAc). Corresponding to previous findings, Mg^2+^ strongly affected the WTA glycosylation profile in WT bacteria, with a significant decrease of 4461 (α1,4-GlcNAc) Fab binding and concomitant increase in 4497 (β1,4-GlcNAc) Fab binding compared to growth in regular TSB (Figure 4A). In contrast, addition of Na^+^ only slightly increased levels of β1,4-GlcNAc, but had no effect on α1,4-GlcNAc levels (Figure 4A). Interestingly, no differences in α1,4- and β1,4-GlcNAc levels were observed in Δ*arlRS* when grown in TSB supplemented with Mg^2+^ or Na^+^ compared to growth in TSB only. We next assessed whether the Mg^2+^-induced WTA glycoswitch was caused by changes in *tarM* promoter activity using our sGFP transcriptional reporter system. Mg^2+^ strongly reduced *tarM* promoter activity in WT but not in the Δ*arlRS* mutant (Figure 4B). These results suggest that Mg^2+^, but not Na+, signals through ArlRS to repress *tarM* promoter activity, which results in increased β1,4-GlcNAc levels.

**Figure 4:**
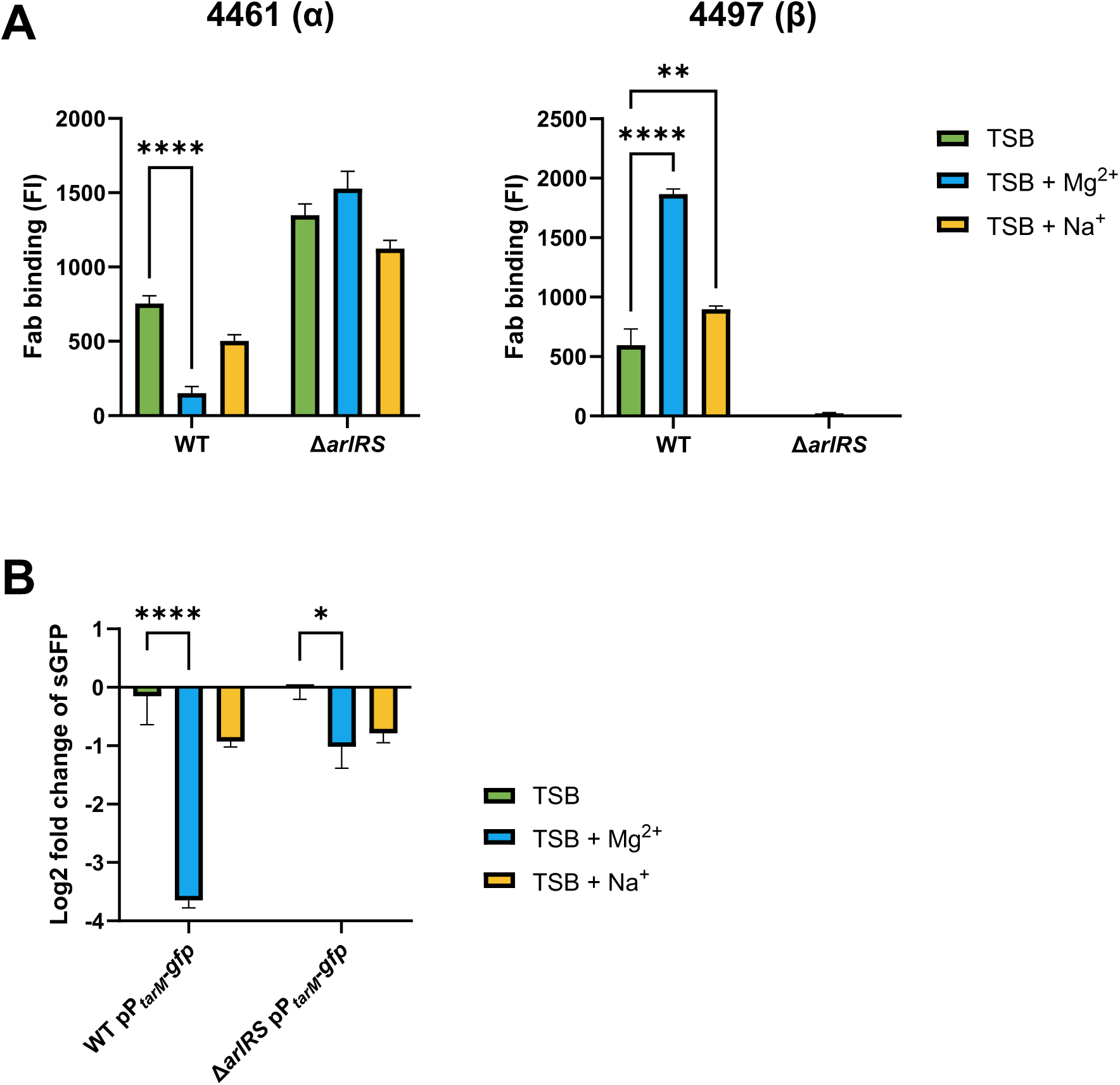
Mg^2+^ drives WTA glycoswitch towards β1,4-GlcNAc and is ArlRS dependent. (A) Levels of WTA α1,4- and β1,4-GlcNAc, quantified by binding of Fab fragments 4461 (α) and 4497 (β) to WT and Δ*arlRS* when grown in TSB or TSB supplemented with 200 mM Mg^2+^ or 200 mM Na^+^. Data is represented as fluorescence intensity (FI) as analyzed by flow cytometry. (B) Promoter activity of *tarM*, displayed as log2 fold change in fluorescence intensity of sGFP in WT pP*_tarM_*-*gfp* or Δ*arlRS* pP*_tarM_*-*gfp* bacteria grown in TSB supplemented with 200 mM Mg^2+^ or 200 mM Na^+^ compared to regular TSB. Data represent three biological replicates ± SEM. Statistical significance was determined using two-way ANOVA with Bonferroni statistical hypothesis testing to correct for multiple comparisons. The mean of each column was compared to the mean of TSB marking only significant comparisons. \**p* < 0.05, \*\**p* < 0.01, \*\*\*\**p* < 0.0001.

### TCS ArlRS represses *tarM* expression through MgrA

In previous research, *tarM* was not identified as part of the ArlRS regulon [41]. However, regulation could be indirect, since ArlRS tightly regulates the expression of the global transcriptional regulators *mgrA* and *spx* by binding to a sequence motif in the promoter region. In turn, MgrA and Spx control the expression of numerous downstream genes [33, 41]. To investigate the link between ArlRS and *tarM* regulation, we analyzed the activity of the *spx* and *mgrA* promoters with the sGFP-reporter system in both WT and the Δ*arlRS* mutant strain. sGFP fluorescence was measured after overnight culture in TSB containing 50, 100, or 200 mM of Mg^2+^ or Na^+^ and was compared to fluorescence of bacteria grown in TSB only. We observed a dose-dependent increase in *mgrA* promoter activity in response to Mg^2+^, but not Na^+^ (Figure 5A). In contrast, the *spx* promoter did not respond to the addition of Mg^2+^ or Na^+^ in the medium (Figure 5A).

**Figure 5:**
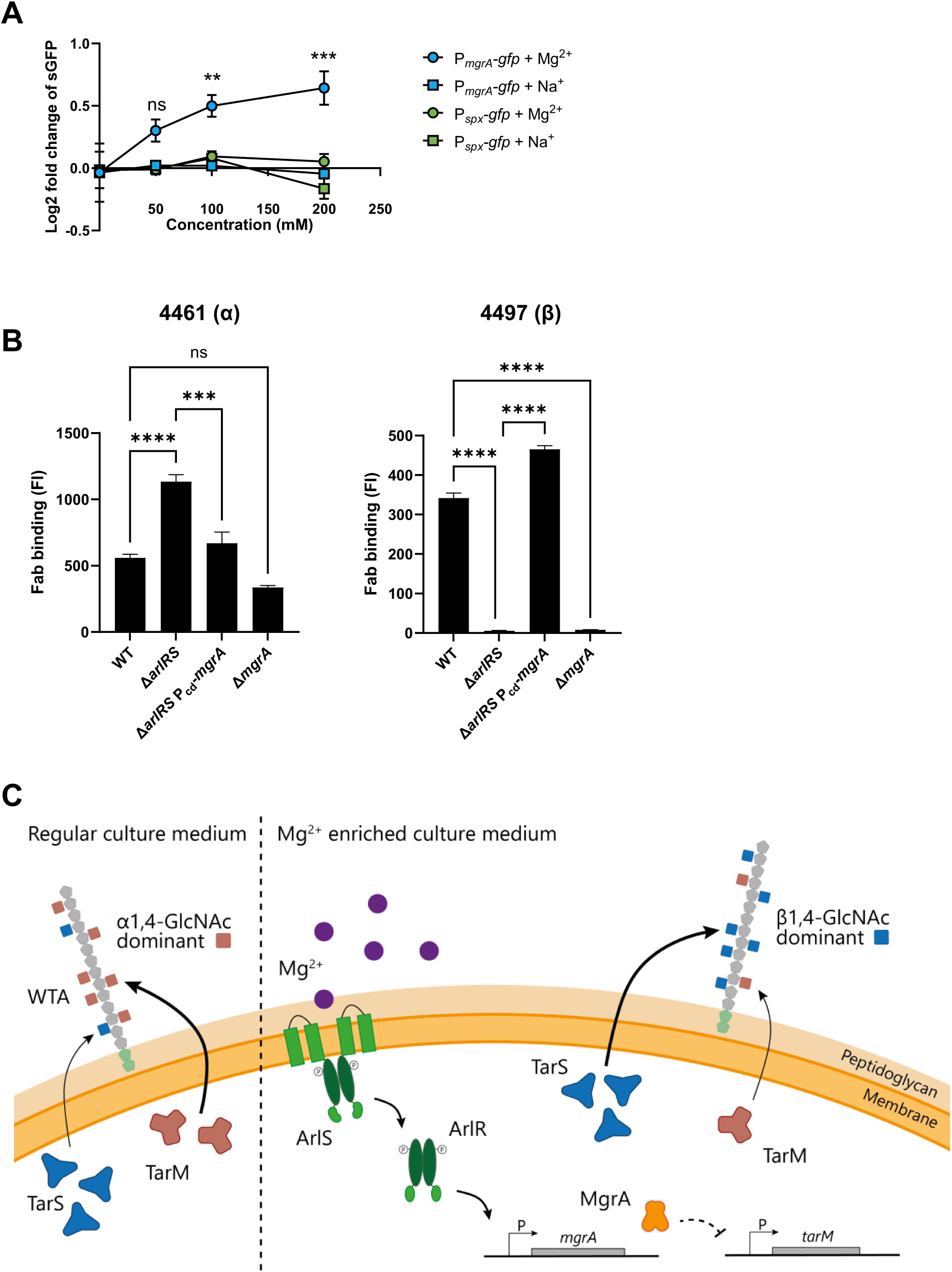
ArlRS represses *tarM* expression through MgrA. (A) Promoter activity of *mgrA* and *spx*, displayed as log2 fold change in fluorescence intensity of sGFP in WT pP*_mgrA_*-*gfp* or WT pP*_spx_*-*gfp* bacteria grown in TSB supplemented with Mg^2+^ or Na^+^ at various concentrations. (B) Levels of α1,4- and β1,4-GlcNAc, quantified by binding of Fab fragments 4461 (α) and 4497 (β) to WT, Δ*arlRS*, the complemented strain Δ*arlRS* PCd-*mgrA* and Δ*mgrA* of *S. aureus* MW2. Data is represented as fluorescence intensity (FI) as analyzed by flow cytometry. Data represent three biological replicates ± SEM. Statistical significance was determined using one-(B) and two-way (A) ANOVA with Bonferroni statistical hypothesis testing to correct for multiple comparisons. \*\**p* < 0.01, \*\*\**p* < 0.001, \*\*\*\**p* < 0.0001, ns=not significant. (C) Schematic overview of the proposed pathway for ArlRS-regulated WTA glycoswitching. Mg^2+^ activates ArlRS, inducing expression of *mgrA*. MgrA in turn either directly or indirectly represses *tarM* expression, allowing TarS to decorate WTA with predominantly β1,4-GlcNAc.

To confirm that MgrA is part of the regulatory network of ArlRS-dependent WTA glycosylation, the native *mgrA* promoter was exchanged with a cadmium-inducible promoter in the Δ*arlRS* mutant (Δ*arlRS* P_Cd_*mgrA*). Importantly, the Δ*arlRS* P_Cd_*mgrA* strain expressed *mgrA* and *tarM* mRNA at levels comparable to those in WT and Δ*arlRS* p*arlRS* (Figure S1). Chromosomal exchange of P_Cd_*mgrA* restored WTA α1,4-GlcNAc and β1,4-GlcNAc glycosylation to WT levels (Figure 5B). Remarkably, while the Δ*mgrA* mutant showed loss of β1,4-GlcNAc, it maintained normal levels of α1,4-GlcNAc compared to WT (Figure 5B). Collectively, these results indicate that ArlRS regulates WTA glycoswitching through the transcription factor MgrA, likely through repression of *tarM* transcription. The proposed pathway is visualized in Figure 5C.

### ArlRS affects β1,4-GlcNAc-dependent functions as langerin binding and phage infection

To assess the functional consequences of ArlRS-dependent glycoswitching, we explored the effects on β1,4-GlcNAc-mediated processes, i.e. langerin binding and phage susceptibility. We have previously shown that langerin, a C-type lectin receptor exclusive to Langerhans cells, recognizes *S. aureus* WTA β1,4-GlcNAc, which subsequently initiates an inflammatory response in the skin [11]. We hypothesized that ArlRS could affect langerin binding, thereby impacting the first response of Langerhans cells. Using recombinant FITC-labeled constructs of the extracellular domains of langerin, we observed that langerin binding was greatly diminished in bacteria lacking *arlRS* compared to WT bacteria (Figure 6A). This phenotype could be restored by complementation with plasmid-expressed *arlRS* (Figure 6A). Correspondingly, langerin binding to WT bacteria could be further enhanced by supplementing the culture medium with Mg^2+^, but not Na^+^ (Figure 6B). In line with our hypothesis, langerin binding to the Δ*arlRS* mutant was not affected by the addition of Na^+^ or Mg^2+^.

**Figure 6:**
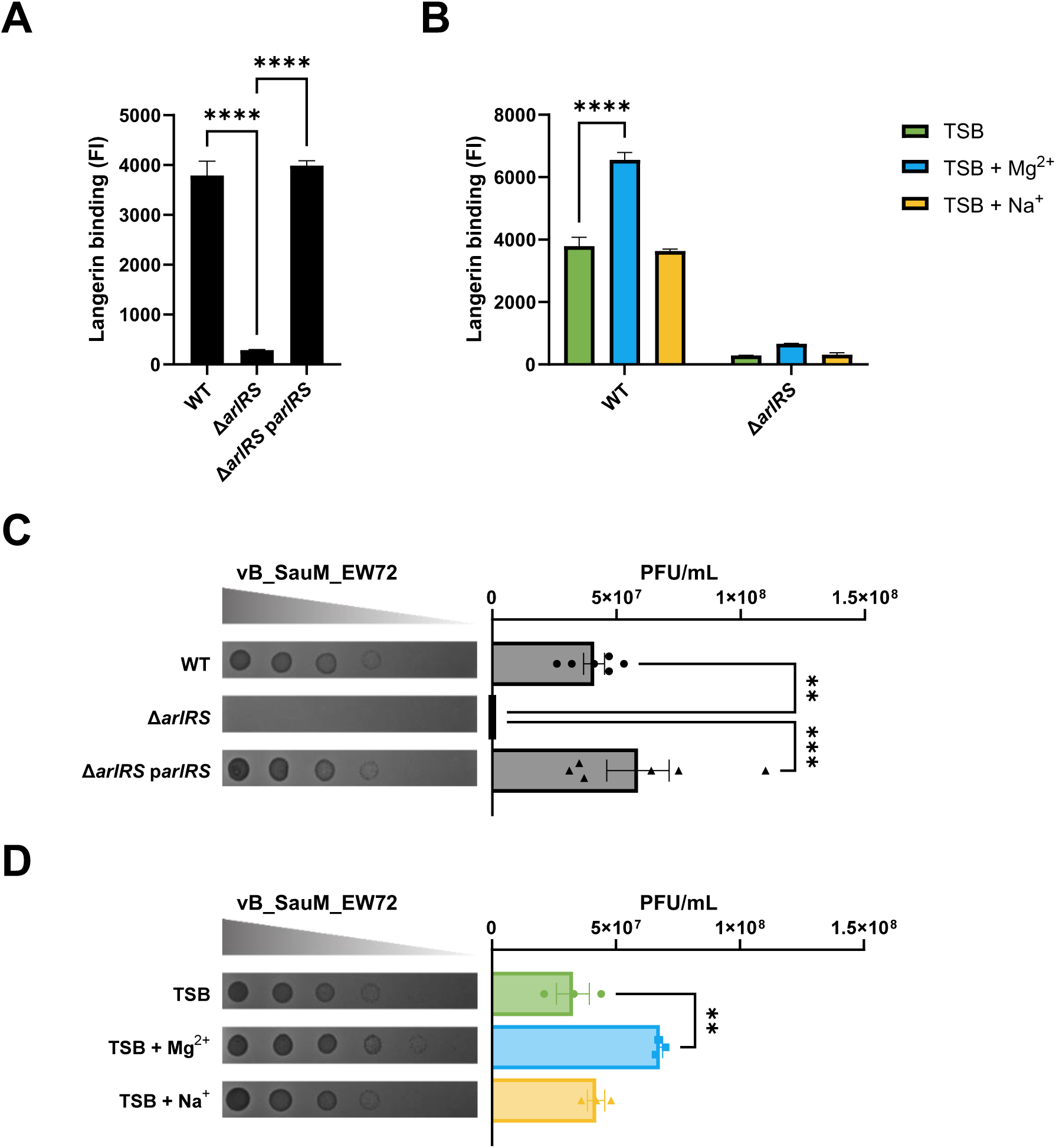
ArlRS activity is required for langerin binding and phage-mediated lysis. Binding of recombinant langerin receptor to overnight cultures of (A) WT, Δ*arlRS* and Δ*arlRS* p*arlRS* bacteria or (B) WT and Δ*arlRS* grown in regular TSB or TSB supplemented with 200 mM Mg^2+^ or Na^+^. Data is represented as fluorescence intensity (FI) as analyzed by flow cytometry from three biological replicates. (C, D) Phage dilutions of vB_SauM_EW72 were spotted on a lawn of (C) WT, Δ*arlRS* or Δ*arlRS* p*arlRS* bacteria or (D) WT bacteria grown in TSB or TSB supplemented with 200 mM Mg^2+^ or Na^+^. Representative images are shown of the formed plaques. PFU/mL was determined from three or six biological replicates. Data are shown as mean ± SEM. Statistical significance was determined using one-(A, C, D) and two-way (B) ANOVA with Bonferroni statistical hypothesis testing to correct for multiple comparisons. For panel B and D, the mean of each column was compared to the mean of TSB marking only significant comparisons. \*\*\**p* < 0.001, \*\*\*\**p* < 0.0001.

WTA glycosylation also affects phage infection as some phages, e.g. Stab20 and Stab20-like phages such as vB_SauM_EW72 [22], require WTA β-GlcNAc moieties to bind to and subsequently infect *S. aureus* [16, 17]. We therefore tested phage susceptibility of our strains. As shown previously [22], we observed phage infection of WT bacteria, with clearly visible plaques that averaged 4*10^7^ PFU/mL (Figure 6C, Figure S2). However, no plaques were observed in the Δ*arlRS* mutant, indicating a 7-log difference in infectivity (Figure 6C). Supplementation with Mg^2+^, but not Na^+^, enhanced infection by some phages in WT bacteria, although this effect was relatively minor (Figure 6D, Figure S3). Together, the langerin binding and phage infection data indicate that ArlRS-dependent glycoswitching also has biologically relevant consequences in host and phage interaction.

## Discussion

In this study, we have shown that the TCS ArlRS, through the transcriptional regulator MgrA, plays an essential role in regulating the specific GlcNAc decoration of the cell wall glycopolymer WTA, affecting phage infection and langerin binding. In one-third of *S. aureus* isolates, WTA is not only decorated with β1,4-GlcNAc through the enzyme TarS, but is co-decorated with α1,4-GlcNAc through the accessory enzyme TarM [12]. These TarM/TarS-positive bacteria can thereby regulate WTA glycosylation to switch between α1,4- or β1,4-GlcNAc-dominant profiles, attuning them to the environmental conditions. The capacity to dynamically regulate the WTA glycoprofile potentially provides a significant survival advantage to *S. aureus* in specific host niches. Moreover, since WTA is a promising target for various alternative treatment options such as vaccines, antibody-dependent therapies and phage therapy, it is important to understand how WTA glycoswitching is regulated to align antimicrobial strategies with WTA glycosylation profiles during infection.

By screening 1,920 transposon mutants, we identified 230 mutants with a divergent α1,4- or β1,4-GlcNAc WTA glycosylation profile. Fifteen of the identified genes had regulatory functions including the three TCS ArlRS, AgrCA and GraRS. Using a panel of isogenic TCS mutants, we confirmed the role of ArlRS in regulating α1,4- and β1,4-GlcNAc WTA glycosylation. Lack of ArlRS resulted in complete loss of β1,4-GlcNAc, with concomitantly increased WTA α1,4-GlcNAc decoration. Moreover, the role of ArlRS in regulating the WTA α1,4/β1,4-GlcNAc ratio was non-redundant, since the WTA glycosylation phenotype could be restored by plasmid-based *arlRS* complementation in a ΔXV mutant, which lacks all 15 non-essential TCS. We were unable to confirm a contribution of *graRS* and *agrCA* in regulating the WTA glycosylation profile, although both TCS have been implicated in the regulation of *tarM* expression [22, 28]. This may be explained by the use of different strain backgrounds, since the NTML is constructed in the *S. aureus* USA300 JE2 background. The TCS deletion mutants used in this study were in *S. aureus* USA400 MW2 and the previous report on the role of AgrCA in the regulation of WTA glycosylation used *S. aureus* Newman [22]. Indeed, strain-specific effects of *agr*-dependent regulation have been previously documented [23]. In addition, glycosylation differences were only observed when stimulating the AgrCA pathway with autoinducing peptides, but not when inactivating this pathway through deletion of *agrA* [22]. Therefore, the effect of AgrCA on WTA glycosylation profiles may depend on the activity and regulation of the quorum-sensing system of the particular strain and the experimental conditions.

Using a *tarM*-specific GFP-promoter and mRNA expression analysis, we showed that ArlRS-mediated regulation of the WTA glycosylation profile resulted from increased expression of *tarM*, without any apparent changes in *tarS* expression. Correspondingly, re-analysis of *S. aureus* transcriptome data generated in 44 *in vitro* and *in vivo* mimicking conditions, demonstrated that *tarM* expression varies substantially, while *tarS* levels remain constant at a relatively high level in all experimental conditions [24, 37]. Regulation of the WTA glycoprofile through regulation of *tarS* may not be possible without affecting general WTA turnover since *tarS* is part of the *tar* WTA biosynthesis operon[24, 37]. Instead, regulation of the accessory *tarM* gene, which is located separated from other *tar* genes, may be better suited to change WTA glycosylation profiles. As a result of increased transcription, the increased levels of TarM likely outcompete TarS at a functional level due to its higher enzymatic activity [42, 43]. *In vitro* biochemical assays with different ratios of recombinant TarM and TarS may be able to confirm this enzymatic competition in the future.

We were able to reduce *tarM* expression and restore WTA β1,4-GlcNAc glycosylation in the Δ*arlRS* mutant by overexpressing *mgrA*, demonstrating that ArlRS-mediated regulation of *tarM* expression requires *mgrA* and not *spx*. MgrA itself was not identified in our screen, since this gene is not included in the NTML. Unexpectedly, deletion of *mgrA* did not completely phenocopy the *arlRS* mutant. Although the *mgrA* mutant showed a complete loss of β1,4-GlcNAc similar to the *arlRS* mutant, *mgrA* deletion did not result in a significant concomitant increase in WTA α1,4-GlcNAc decoration. It is already known that the phenotypes of *arlRS* and the *mgrA* deletion strains show overlap due to overlapping regulons [31, 34, 41, 44], but *tarM* is not part of either regulon [41, 45]. A similar observation is made in a recent paper, in which MgrA is identified as the regulator of the *ssc* genes encoding for a novel *S. aureus* secondary glycopolymer, but no canonical *mgrA* binding site could be found [46]. Alternatively, the deletion of *mgrA* may interfere with other regulatory pathways. Indeed, MgrA represses expression of the transcription factors SarV and AtlR [33], which may indicate that MgrA regulates *tarM* expression indirectly. Therefore, the molecular mechanisms by which MgrA represses *tarM* remain elusive. Furthermore, despite recent advances in RNA sequencing or prediction of transcription factor binding sequences, our observations stress the importance of combining phenotypic screening with experimental work to confirm direct gene regulation [33, 47].

Shifts in WTA glycosylation profiles can be induced by general stress, such as high salt concentrations [21]. The medium previously used contained both high Na^+^ and Mg^2+^ concentrations. We first demonstrated that salt-induced effects on WTA glycoprofiles were induced by high levels of Mg^2+^ but not Na^+^. We next showed that these Mg^2+^-induced changes in the WTA glycoprofile depended on repression of *tarM* promoter activity, which required the ArlRS TCS. Our data extend earlier research on the link between ArlRS, Mg^2+^, and WTA, where it was shown that ArlRS and Mg^2+^ repressed the *dlt* operon [29], which decorates WTA with D-alanine residues. In addition to transcriptional changes, biochemical studies have shown that Mg^2+^ acts as a co-factor to increase TarS enzymatic activity, which in our conditions may have additionally boosted β1,4-GlcNAc modifications [43]. Unexpectedly, we also detected a slight increase in WTA β1,4-GlcNAc levels when TSB was supplemented with Na^+^. However, Na^+^ did not affect *tarM* or *mgrA* promoter activity. Therefore, these Na^+^-induced effects most likely involved a different pathway independent of ArlRS or MgrA.

Mg^2+^ is an essential mineral for many basic biological processes and one of the four most common cations in the human body [36]. Ninety-nine percent of the total amount of Mg^2+^ is found in bones, muscle and soft tissue [48]. The remaining Mg^2+^ is present in red blood cells and serum [48, 49]. *S. aureus* can naturally infect the skeletal organ [50]. This Mg^2+^-enriched niche seems to favor chronic infections and to induce biofilm formation of *S. aureus* [51, 52]. Although in our study Mg^2+^ clearly had a great influence on *S. aureus*, there are no studies yet that analyzed the complete gene set regulated by Mg^2+^, as has been done for *Streptococcus pyogenes* [53]. In *S. pyogenes*, CovS of the streptococcal TCS CovRS has been identified as the sensor for Mg^2+^ and has 32% sequence identity with ArlS. As no dedicated activation signal for ArlRS has been identified, we propose that Mg^2+^ may be a primary stressor for the ArlRS TCS in *S. aureus*. In line with this suggestion, we identified two metal transporters, MntB (manganese) and MgtE (magnesium), in the 4461 (α1,4-GlcNAc) screen, of which both metals have previously been implicated with ArlRS [29, 35]. These findings suggest an important role of metal ion availability in the *S. aureus* WTA glycoswitch. Moreover, it highlights how *S. aureus* could undergo phenotypic WTA modifications in environments with high Mg^2+^ concentrations such as bone, muscle and soft tissue through the sensing and transcriptional regulation of ArlRS and MgrA.

ArlRS plays a role in several host-pathogen interactions [30–35], including in several animal models [31, 33, 34, 44]. To probe the potential functional consequences of ArlRS-mediated WTA glycoswitching for biologically-relevant interactions, we performed studies to investigate interaction with human langerin and the impact on infection by β1,4-GlcNAc-dependent phages. Langerin is a pattern recognition receptor unique to innate Langerhans cells that specifically binds staphylococcal WTA β-GlcNAc and is hindered by co-presence of α1,4-GlcNAc. This interaction acts as first response in the defense against *S. aureus* in the skin through the induction of skin inflammation specifically neutrophil recruitment [11]. As expected, the increased β1,4-GlcNAc levels in response to high Mg^2+^ and dependence on ArlRS activation resulted in increased Langerin binding. How the conditions in human skin including the concentration of Mg^2+^ would affect ArlRS activation and subsequent WTA glycosylation profiles is currently unknown but warrants further investigation.

With regard to phage infection, the effect of WTA glycoswitching is a double-edged sword. Whereas bacteria may profit from phages to gain favorable genes via horizontal gene transfer [4], lytic phages can also kill bacteria [54]. WTA decoration with GlcNAc is required for successful infection by siphophages, podophages and some myophages [5, 15–17], but some phages require GlcNAc to be present in a specific configuration [55]. In this study, all used phages depend on the WTA β1,4-GlcNAc modification to infect and lyse *S. aureus* [22]. Consequently, it was expected that phage infection was completely abolished in the Δ*arlRS* mutant lacking β1,4-GlcNAc. This is consistent with a previous observation that after co-culture, *S. aureus* bacteria resistant to the β1,4-GlcNAc-dependent podophage carried an inactivating mutation in ArlRS [56]. In contrast to our finding with langerin and Fab binding experiments, the influence of Mg^2+^ on phage infection was modest. This may be due to the long infection period on the agar plates that did not contain extra Mg^2+^. Nevertheless, our results still suggest that Mg^2+^ concentrations influence *S. aureus* phage infection. This is of importance for choosing the right phage therapy, as an isolate may be different with regard to WTA glycosylation depending on the location of the infection, such as a osteomyelitis versus endocarditis.

In summary, we identified the critical importance of ArlRS in the dynamic regulation of the WTA GlcNAc decoration in *S. aureus*. ArlRS was activated by high concentrations of Mg^2+^ and subsequently activated MgrA and upregulated *tarM* expression, resulting in WTA glycoswitching from a β1,4-GlcNAc-dominated to an α1,4-GlcNAc-dominated profile. As WTA glycosylation patterns impact many biologically- and infection-relevant interactions, more research is needed to determine differential GlcNAc levels of *S. aureus* isolates in specific tissues such as bone and skin, common sites of (chronic) *S. aureus* infections. This is especially important since *in vitro* testing of GlcNAc levels in clinical isolates after routine culture or screening for gene presence do not correlate with *in vivo* WTA glycosylation patterns. These explorations could help prioritize the design of new antimicrobial strategies such as vaccines and phage therapy targeting specific WTA glycan modifications.

## Acknowledgements

We thank S. Man-Bovenkerk for technical assistance and A.R. Temming for providing the Fab clones 4461 and 4497. This work was supported by the Vici (09150181910001) research program to N.M.v.S., which is financed by the Dutch Health Council (NWO) and by the NNF foundation, NNF22OC0077593 to E.L. and H.I. and Danmarks Frie Forskningsfond, 2035-00110B to H.I..

## Materials & Methods

### Bacterial strains and culture conditions

All *S. aureus* cultures were grown in Tryptic Soy Broth (TSB, Oxoid) overnight at 37°C with continuous shaking. The Nebraska Transposon Mutant Library (NTML; 1,920 arrayed mutants of *S. aureus* strain JE2, harboring both *tarM* and *tarS*) [38] was grown in presence of 5 μg/mL erythromycin (Sigma). *S. aureus* complemented strains were grown with addition of 10 μg/mL erythromycin or 10 μg/mL chloramphenicol (Sigma), depending on the resistance marker used. When required, TSB was enriched with MgCl_2_ (TSB+Mg^2+^, Merck) or NaCl (TSB+Na^+^, Merck) in a concentration of 200 mM unless stated otherwise. *Escherichia coli* strain DC10B [57] was grown at 37°C with continuous shaking in Lysogeny broth (LB) supplemented with 100 μg/mL ampicillin (Sigma) when appropriate. All strains used in this study are listed in Table S1.

### Analysis of *tarM* and *tarS* mRNA transcription levels

The transcriptome of *S. aureus* HG001 grown in 44 different experimental conditions was analyzed in earlier research [24]. The transcript levels of *tarS* (SAOUHSC_00228) and *tarM* (SAOUHSC_00973) were downloaded from the AureoWiki repository [37]. Averages of biological replicates were calculated and plotted, comparing *tarM* and *tarS* expression in all different growth conditions. Statistical difference in variance of the transcript levels was calculated using an F test with GraphPad Prism 10.2.0.

### NTML screen for WTA α1,4- and β1,4-*N*-acetylglucosamine glycosylation levels

The protocol for immunoblotting was based on Weber *et al*. [58] with some modifications. Each of the 1,920 NTML [38] mutants was inoculated from frozen stock into a single well of a 96-well round bottom plate (Sigma) containing TSB with 5 μg/mL erythromycin. The next day, 2 μl of the overnight culture was spotted in duplicate onto Tryptic Soy Agar (TSA, Oxoid) plates containing 5 μg/mL erythromycin. Plates were incubated overnight and bacteria were transferred from each plate to a 0.45 μm nitrocellulose membrane (Biorad). Each membrane was dried for 20 min and subsequently washed with demineralized water and Tris-Buffered Saline (TBS). The membranes were blocked for one hour with TBS supplemented with 1% Tween20 (TBST, Fisher Scientific) and 5% bovine serum albumin (BSA; Sigma) and washed with TBST. The glycosylation of each individual transposon mutant was analyzed using the specific Fab clones 4461 (α1,4-GlcNAc) and 4497 (β1,4-GlcNAc) [59] by incubating the membranes for 1 hour with either 0.1 μg/mL 4461 or 4497 Fabs in TBST. After washing with TBST, the membranes were incubated for 1 hour with a 1:1,000 dilution of Goat Fab Anti-Human IgG Kappa-AP (Southern Biotech) and washed again with TBST. Fab staining, representing WTA glycosylation, was visualized using Vector blue AP Substrate (Vector laboratory). The reaction was stopped with a quick wash with demineralized water and blots were imaged using a Uvitec Platinum V10 imager. Differential glycosylation patterns were assessed by visual comparison of the overall staining intensity of mutants on a single blot.

### Bacterial cloning

MW2 wild type (WT) bacteria and a panel of cognate TCS deletion mutants were previously generated [25, 44]. Complementation mutants of ΔXV and Δ*arlRS* were constructed using the pCN51 plasmid under control of a cadmium promoter, yielding ΔXV p*arlRS* and Δ*arlRS* p*arlRS* [25]. Cadmium was not used for the expression of genes as the leaky expression on this plasmid was sufficient to restore the phenotype [25]. The *arlRS* deletion mutant complemented with *mgrA* (Δ*arlRS* P_Cd_-*mgrA*) was constructed by Burgui *et al*. [44] via allelic exchange of the *mgrA* promoter with the cadmium inducible promoter. As this complementation is chromosomal rather than plasmid-based, 5 μM cadmium was added to culture medium to reach sufficient ArlRS-independent expression of *mgrA*.

### Analysis of α1,4- and β1,4-*N*-acetylglucosamine glycosylation levels

WTA glycosylation with either α1,4- or β1,4-GlcNAc was analyzed using specific Fab fragments. Overnight cultures in TSB with or without appropriate antibiotics were resuspended in PBS + 0.1% BSA at an optical density 600 nm (OD_600_) of 0.4 and incubated with 3.3 μg/mL Fab fragments 4461 (α1,4-GlcNAc) or 4497 (β1,4-GlcNAc) [59]. After washing, bacteria were incubated with 1:200 goat Fab human Kappa IgG Alexa Fluor 647 (Southern Biotech). Bacteria were washed and fixed in 1% paraformaldehyde (PFA, Sigma-Aldrich), and analyzed by flow cytometry (BD FACSCanto II Flow Cytometer, BD Bioscience) using FlowJo version 10 (BD Bioscience).

### Promoter activity analysis using *promoter*::*gfp* translational fusions

Promoter activity of *tarM*, *mgrA,* and *spx* was analyzed using a fluorescent reporter system with the plasmid pCM29, which contains a *SarA*-P1 promoter upstream of sGFP [40]. Specific promoter regions of *mgrA*, *spx* [33, 41], and *tarM* [60] were cloned into pCM29 via restriction enzyme digest to replace the *SarA*-P1 promoter. Resulting plasmids were first transformed in *E. coli* DC10B and subsequently transformed into MW2 WT (resulting in WT P*_mgrA_-gfp*, WT P*_spx_-gfp*, and WT P*_tarM_-gfp*) and in Δ*arlRS* (resulting in Δ*arlRS* P*_mgrA_-gfp*, Δ*arlRS* P*_spx_-gfp*, and Δ*arlRS* P*_tarM_-gfp*). Used primers are listed in Table S2.

sGFP-reporter strains were grown overnight in TSB supplemented with 10 μg/mL chloramphenicol and Mg^2+^ or Na^+^ if appropriate. Cultures were adjusted to an OD_600_ of 0.4 in PBS + 0.1% BSA and fixed with 1% PFA (Sigma-Aldrich). sGFP intensity was analyzed by flow cytometry (BD FACSCanto II Flow Cytometer, BD Bioscience) using FlowJo version 10 (BD Bioscience).

### Absolute quantification of mRNA transcripts by reverse transcriptase (RT)-qPCR

As ArlRS has an extensive influence on expression levels of many genes, we quantified the absolute number of mRNA copies for *tarM* and *mgrA* to circumvent the use of housekeeping genes. Briefly, pellets of overnight bacterial cultures were resuspended in TRIzol reagent (Invitrogen) and bacterial cells were disrupted using 0.1 mm zirconium sand and 2 mm glass beads using a Magna-lyser for 2 min at 6,000 rpm. Total RNA was extracted using Direct-zol RNA miniprep-kit (Zymo Research) according to manufacturer’s protocol followed by a DNase treatment using the Turbo DNA-free kit (Invitrogen). cDNA was synthesized using the Maxima H Minus cDNA Synthesis Master Mix (ThermoFisher) and absence of genomic DNA was confirmed through no-RT PCRs. RT-qPCR was performed using a standard curve as described by Goerke *et al.* [61]. In short, the genes were amplified with regular PCR using genomic DNA of *S. aureus* MW2 as template and primers listed in Table S2. The amplicons were visualized on gel for specificity, cleaned using GeneJET PCR Purification Kit (Thermo Scientific) and DNA concentration was measured with the Qubit 1X dsDNA HS Assay Kit (Invitrogen) as analyzed with a Qubit 4 Fluorometer. The sequence-specific standard curves of *tarM*, *tarS* and *mgrA* were then generated using a 10-fold serial dilution (10^2^ to 10^8^ copies/reaction). Quantitative RT-PCR was performed in triplicate on the standard curve and cDNA samples using the PowerTrack SYBR Green Master Mix (Thermo Scientific) in the CFX384 RT-PCR instrument (Bio-Rad) with CFX Maestro 5.0 software. Copy numbers per reaction of samples were determined via inter- or extrapolation of the Ct values of cDNA samples.

### Langerin binding

*S. aureus* langerin binding was analyzed as described previously [11, 59]. Recombinant FITC-labelled construct of human langerin was kindly provided by Prof. C. Rademacher, University of Vienna, Vienna, Austria [62]. In short, overnight cultures were resuspended in TSM buffer [2.4 g/L Tris (Sigma-Aldrich), 8.77 g/L, NaCl (Merck), 294 mg/L CaCl_2_(H_2_O)_2_ (Merck), 294 mg/L MgCl_2_(H_2_O)_6_ (Merck), containing 0.1% BSA, pH 7] at an OD_600_ of 0.4 and incubated with 20 μg/mL recombinant langerin. Langerin binding was analyzed by flow cytometry (BD FACSCanto II Flow Cytometer, BD Bioscience).

### Phage spot assay

All phages used in this study are listed in Table S1. Phage susceptibility of *S. aureus* was tested via phage spot assay [22] using the double agar overlay method [63]. Briefly, overnight cultures were mixed with phage top agar (20 g/L Nutrient Broth No. 2; 3.5 g/L Agar No.1 supplemented with 10 mM CaCl_2_) and poured over phage base agar (20 g/L of Nutrient Broth No. 2; 7 g/L Agar No.1 supplemented with 10 mM CaCl_2_) to generate indicator plates. Phage lysates were prepared as described earlier [64] and were diluted to 1×10^8^ plaque forming units (PFU)/mL followed by a 10-fold serial dilution in phage buffer (1 mM MgSO_4_, 4 mM CaCl_2_, 50 mM Tris-HCl, 100 mM NaCl, pH 8.0). Respective dilutions were spotted on top of the indicator plates. After overnight incubation at 37°C, plaques were counted, PFU/mL determined and plates were imaged (BioRad ChemiDoc XRS+ imager).

### Statistical analysis

Statistical analysis was performed using GraphPad Prism 10.2.0. The F test was used to compare the variance of two groups, the Student’s *t* test and one- and two-way ANOVA with Bonferroni statistical hypothesis testing to correct for multiple comparisons were used to determine significant difference (*p* < 0.05) between two or more groups. All values are reported as mean with standard error of the mean (SEM) of three biological replicates unless indicated otherwise.

## Supplementary data

**Figure S1.**
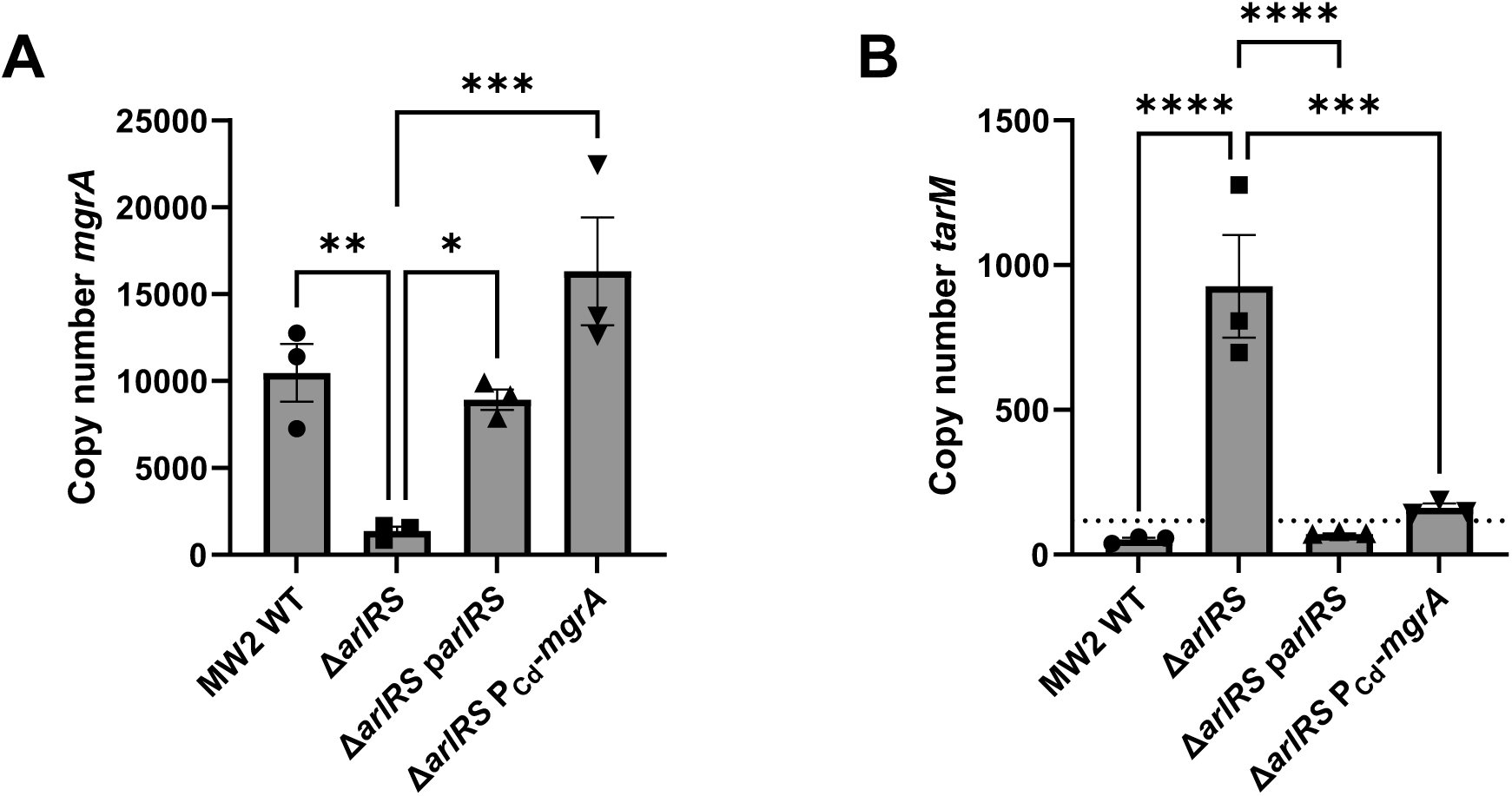
Expression of *mgrA* and *tarM* in Δ*arlRS* and complemented strains. mRNA copy number of (A) *mgrA* and (B) *tarM* as measured with qPCR in WT, Δ*arlRS* and complemented bacteria Δ*arlRS* p*arlRS* and Δ*arlRS* P_Cd_-*mgrA*. Symbols below the dotted line represent extrapolated values. Data represent three biological replicates ± SEM. First three columns of panel B (WT, Δ*arlRS* and Δ*arlRS* p*arlRS*) are the same data as Figure 3C shown in the main article. Statistical significance was determined using one-way ANOVA with Bonferroni statistical hypothesis testing to correct for multiple comparisons.\**p* < 0.05, \*\**p* < 0.01, \*\*\**p* < 0.001, **** *p* < 0.0001.

**Figure S2.**
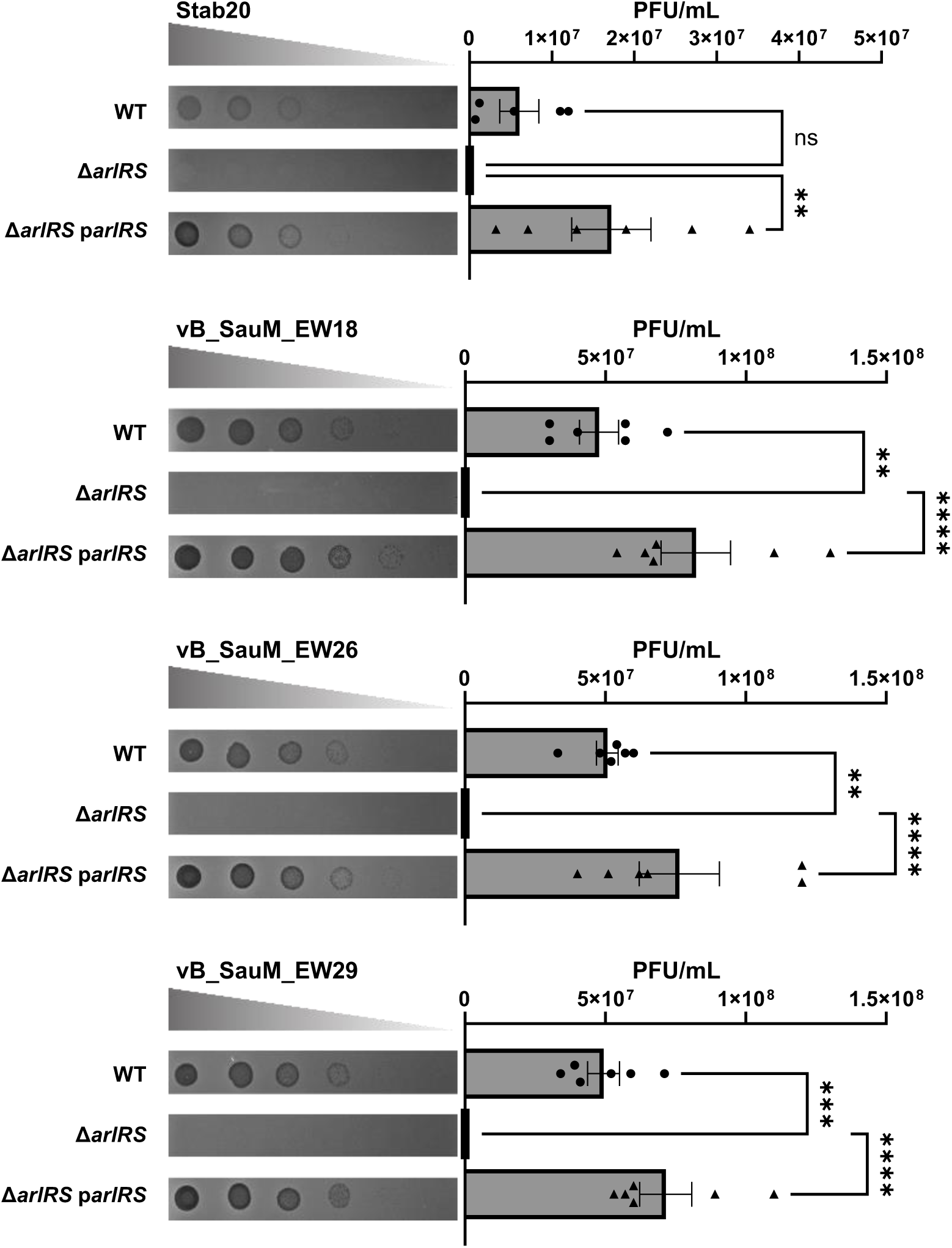
ArlRS is required for successful infection of Stab20 and Stab20-like phages in *S. aureus*. Phage dilutions of Stab20 or the Stab20-like phages vB_SauM_EW18, vB_SauM_EW26, vB_SauM_EW29 were spotted on a lawn of WT, Δ*arlRS* or Δ*arlRS* p*arlRS* bacteria. Representative images are shown of the formed plaques. PFU/mL were counted from six biological replicates. Data are shown as mean ± SEM. Statistical significance was determined using one-way ANOVA with Bonferroni statistical hypothesis testing to correct for multiple comparisons. \*\**p* < 0.01, \*\*\**p* < 0.001, \*\*\*\**p* < 0.0001, ns=not significant.

**Figure S3.**
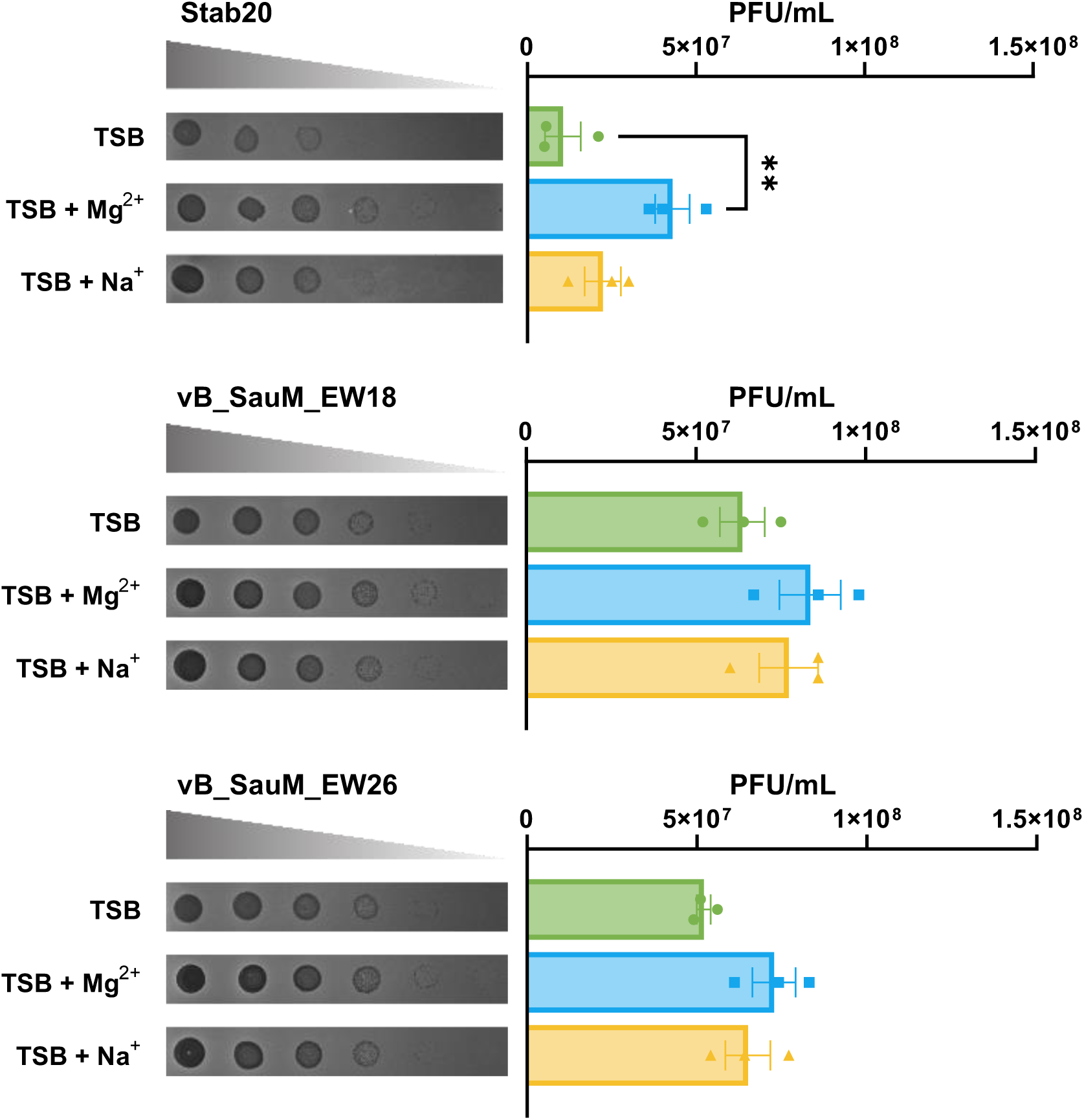
Mg^2+^ enhances infection by Stab20 and Stab20-like phages in *S. aureus*. Phage dilutions of Stab20 or the Stab20-like phages vB_SauM_EW18, vB_SauM_EW26, vB_SauM_EW29 were spotted on a lawn of WT bacteria grown in TSB or TSB supplemented with 200 mM Mg^2+^ or Na^+^. Representative images are shown of the formed plaques. PFU/mL were counted from three biological replicates. Data are shown as mean ± SEM. The mean of each column was compared to the mean of TSB marking only significant comparisons. \**p* < 0.05, ns=not significant.

**Table S1.**
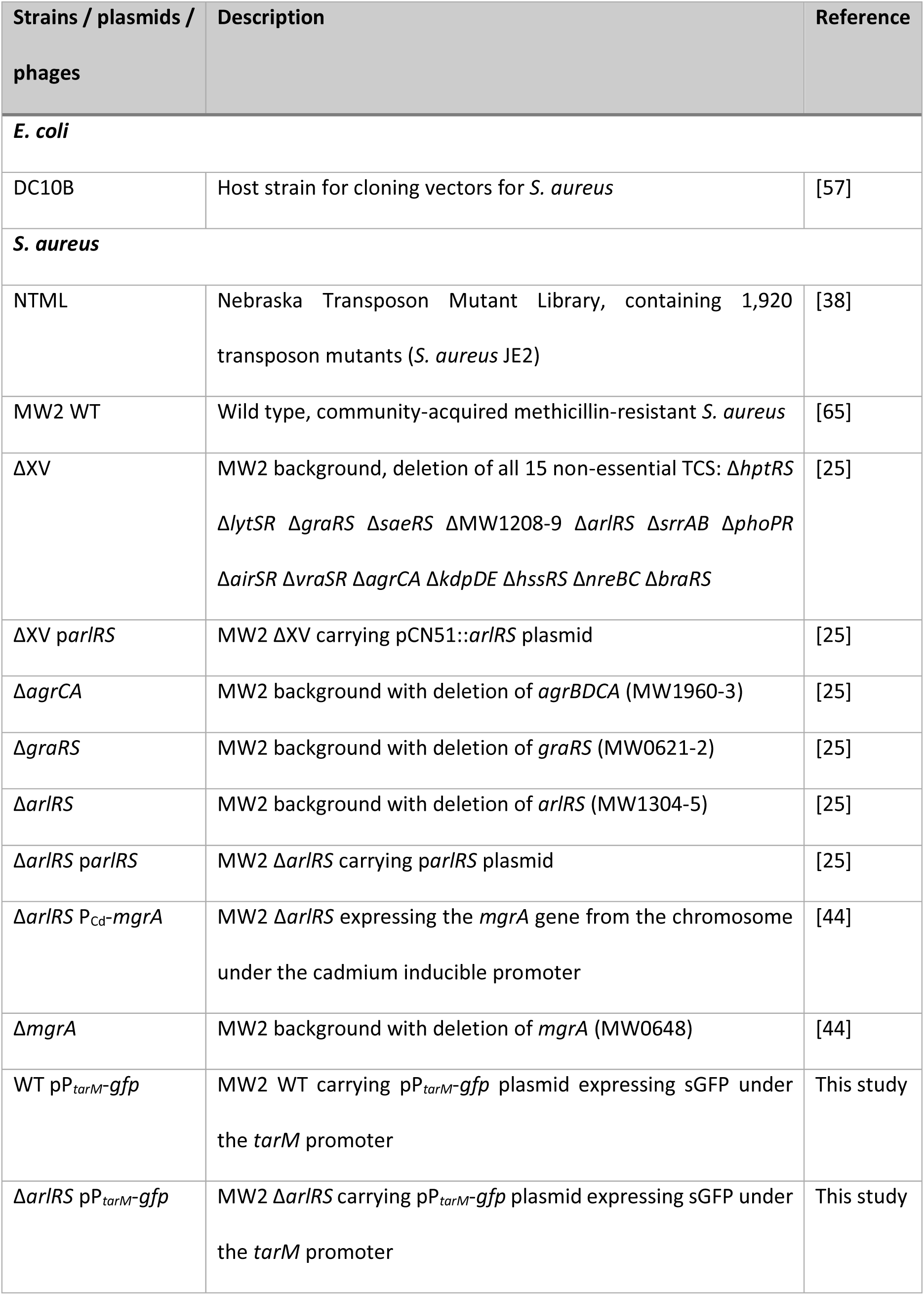
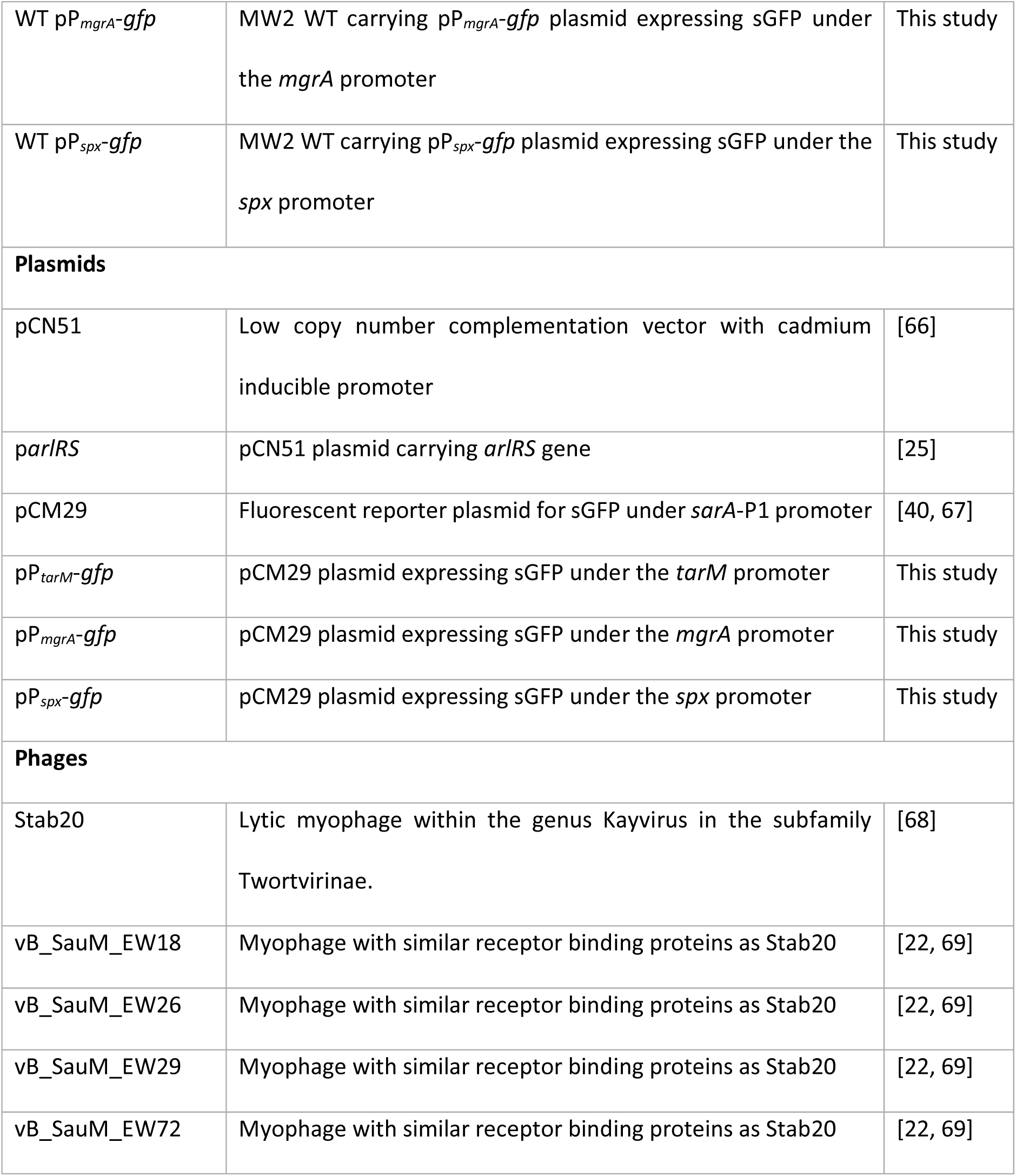
Overview of strains, plasmids and phages used in this study.

**Table S2.**
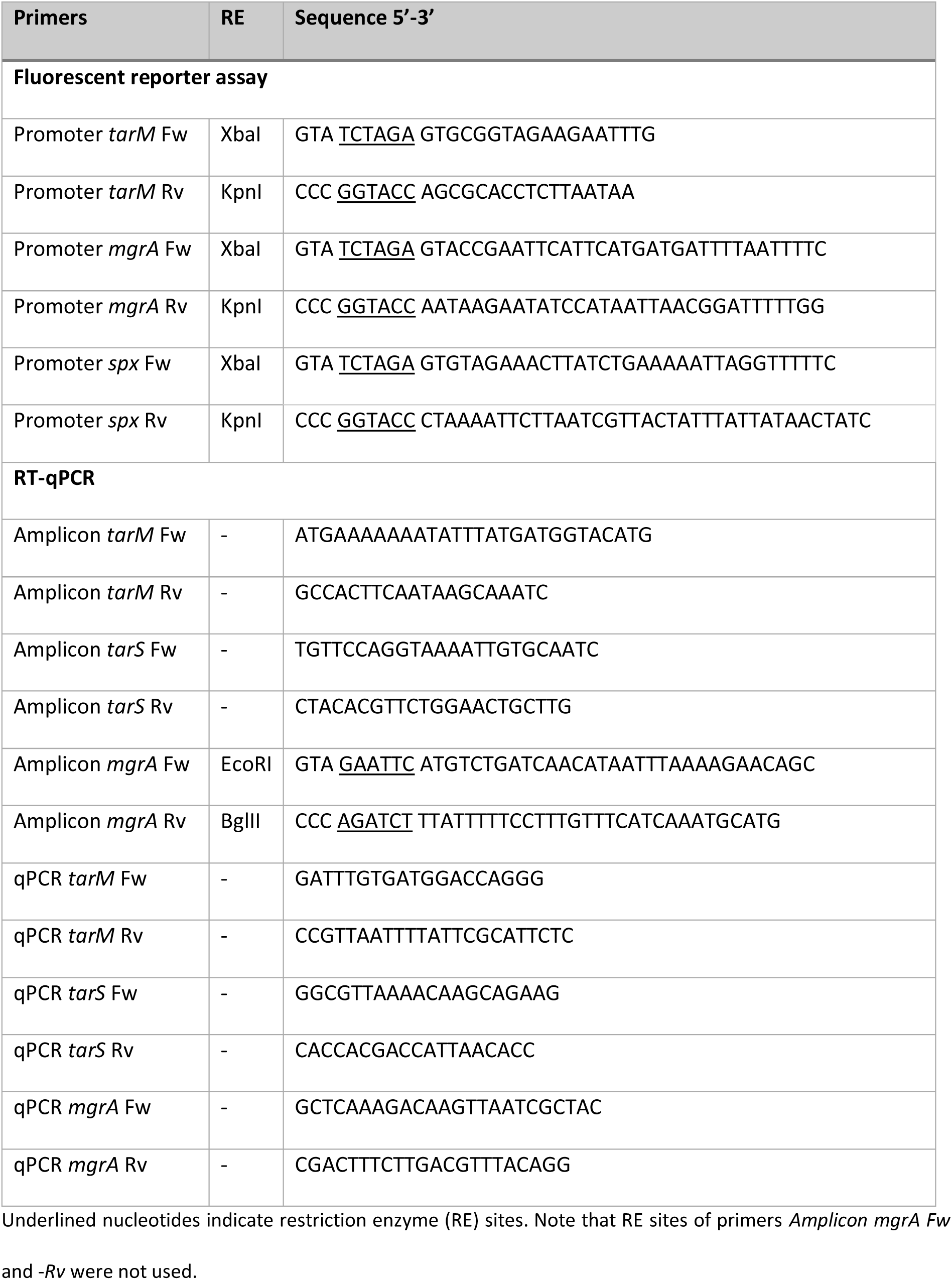
Overview of primers used in this study.

**Table S3: NTML mutants showing aberrant 4461 or 4497 binding levels. See excel file.**

## References

1. Antimicrobial Resistance Collaborators. Global burden of bacterial antimicrobial resistance in 2019: a systematic analysis. Lancet. 2022;399(10325):629–55. doi: 10.1016/S0140-6736(21)02724-0.

2. Tacconelli E, Carrara E, Savoldi A, Harbarth S, Mendelson M, Monnet DL, et al. Discovery, research, and development of new antibiotics: the WHO priority list of antibiotic-resistant bacteria and tuberculosis. Lancet Infect Dis. 2018;18(3):318–27. doi: 10.1016/S1473-3099(17)30753-3.

3. Brown S, Xia G, Luhachack LG, Campbell J, Meredith TC, Chen C, et al. Methicillin resistance in *Staphylococcus aureus* requires glycosylated wall teichoic acids. Proceedings of the National Academy of Sciences of the United States of America. 2012;109(46):18909–14. doi: 10.1073/pnas.1209126109.

4. Winstel V, Liang C, Sanchez-Carballo P, Steglich M, Munar M, Broker BM, et al. Wall teichoic acid structure governs horizontal gene transfer between major bacterial pathogens. Nat Commun. 2013;4(1):2345. doi: 10.1038/ncomms3345.

5. Xia G, Corrigan RM, Winstel V, Goerke C, Grundling A, Peschel A. Wall teichoic Acid-dependent adsorption of staphylococcal siphovirus and myovirus. J Bacteriol. 2011;193(15):4006–9. doi: 10.1128/JB.01412-10.

6. Winstel V, Kuhner P, Salomon F, Larsen J, Skov R, Hoffmann W, et al. Wall Teichoic Acid Glycosylation Governs *Staphylococcus aureus* Nasal Colonization. mBio. 2015;6(4):e00632. doi: 10.1128/mBio.00632-15.

7. Baur S, Rautenberg M, Faulstich M, Grau T, Severin Y, Unger C, et al. A nasal epithelial receptor for *Staphylococcus aureus* WTA governs adhesion to epithelial cells and modulates nasal colonization. PLoS Pathog. 2014;10(5):e1004089. doi: 10.1371/journal.ppat.1004089.

8. Lehar SM, Pillow T, Xu M, Staben L, Kajihara KK, Vandlen R, et al. Novel antibody-antibiotic conjugate eliminates intracellular *S. aureus*. Nature. 2015;527(7578):323–8. doi: 10.1038/nature16057.

9. Jung DJ, An JH, Kurokawa K, Jung YC, Kim MJ, Aoyagi Y, et al. Specific serum Ig recognizing staphylococcal wall teichoic acid induces complement-mediated opsonophagocytosis against *Staphylococcus aureus*. J Immunol. 2012;189(10):4951–9. doi: 10.4049/jimmunol.1201294.

10. van Dalen R, Molendijk MM, Ali S, van Kessel KPM, Aerts P, van Strijp JAG, et al. Do not discard *Staphylococcus aureus* WTA as a vaccine antigen. Nature. 2019;572(7767):E1–E2. doi: 10.1038/s41586-019-1416-8.

11. van Dalen R, De La Cruz Diaz JS, Rumpret M, Fuchsberger FF, van Teijlingen NH, Hanske J, et al. Langerhans Cells Sense *Staphylococcus aureus* Wall Teichoic Acid through Langerin To Induce Inflammatory Responses. mBio. 2019;10(3). doi: 10.1128/mBio.00330-19.

12. Tamminga SM, Volpel SL, Schipper K, Stehle T, Pannekoek Y, van Sorge NM. Genetic diversity of *Staphylococcus aureus* wall teichoic acid glycosyltransferases affects immune recognition. Microb Genom. 2022;8(12):2022.05.25.492469. doi: 10.1099/mgen.0.000902.

13. Xia G, Maier L, Sanchez-Carballo P, Li M, Otto M, Holst O, et al. Glycosylation of wall teichoic acid in *Staphylococcus aureus* by TarM. J Biol Chem. 2010;285(18):13405–15. doi: 10.1074/jbc.M109.096172.

14. Winstel V, Xia G, Peschel A. Pathways and roles of wall teichoic acid glycosylation in *Staphylococcus aureus*. Int J Med Microbiol. 2014;304(3-4):215–21. doi: 10.1016/j.ijmm.2013.10.009.

15. Li X, Gerlach D, Du X, Larsen J, Stegger M, Kühner P, et al. An accessory wall teichoic acid glycosyltransferase protects *Staphylococcus aureus* from the lytic activity of Podoviridae. Sci Rep. 2015;5(1):17219. doi: 10.1038/srep17219.

16. Azam AH, Hoshiga F, Takeuchi I, Miyanaga K, Tanji Y. Analysis of phage resistance in *Staphylococcus aureus* SA003 reveals different binding mechanisms for the closely related Twort-like phages ɸSA012 and ɸSA039. Appl Microbiol Biotechnol. 2018;102(20):8963–77. doi: 10.1007/s00253-018-9269-x.

17. Estrella LA, Quinones J, Henry M, Hannah RM, Pope RK, Hamilton T, et al. Characterization of novel *Staphylococcus aureus* lytic phage and defining their combinatorial virulence using the OmniLog(R) system. Bacteriophage. 2016;6(3):e1219440. doi: 10.1080/21597081.2016.1219440.

18. Takahashi K, Kurokawa K, Moyo P, Jung DJ, An JH, Chigweshe L, et al. Intradermal immunization with wall teichoic acid (WTA) elicits and augments an anti-WTA IgG response that protects mice from methicillin-resistant *Staphylococcus aureus* infection independent of mannose-binding lectin status. PLoS One. 2013;8(8):e69739. doi: 10.1371/journal.pone.0069739.

19. Howden BP, Giulieri SG, Wong Fok Lung T, Baines SL, Sharkey LK, Lee JYH, et al. *Staphylococcus aureus* host interactions and adaptation. Nature Reviews Microbiology. 2023;21(6):380–95. doi: 10.1038/s41579-023-00852-y.

20. van Dalen R, Peschel A, van Sorge NM. Wall Teichoic Acid in *Staphylococcus aureus* Host Interaction. Trends Microbiol. 2020;28(12):985–98. doi: 10.1016/j.tim.2020.05.017.

21. Mistretta N, Brossaud M, Telles F, Sanchez V, Talaga P, Rokbi B. Glycosylation of *Staphylococcus aureus* cell wall teichoic acid is influenced by environmental conditions. Sci Rep. 2019;9(1):3212. doi: 10.1038/s41598-019-39929-1.

22. Yang J, Bowring JZ, Krusche J, Lehmann E, Bejder BS, Silva SF, et al. Cross-species communication via agr controls phage susceptibility in *Staphylococcus aureus*. Cell Rep. 2023;42(9):113154. doi: 10.1016/j.celrep.2023.113154.

23. Jenul C, Horswill AR. Regulation of *Staphylococcus aureus* Virulence. Microbiol Spectr. 2019;7(2). doi: 10.1128/microbiolspec.GPP3-0031-2018.

24. Mäder U, Nicolas P, Depke M, Pane-Farre J, Debarbouille M, van der Kooi-Pol MM, et al. *Staphylococcus aureus* Transcriptome Architecture: From Laboratory to Infection-Mimicking Conditions. PLoS Genet. 2016;12(4):e1005962. doi: 10.1371/journal.pgen.1005962.

25. Villanueva M, Garcia B, Valle J, Rapun B, Ruiz de Los Mozos I, Solano C, et al. Sensory deprivation in *Staphylococcus aureus*. Nat Commun. 2018;9(1):523. doi: 10.1038/s41467-018-02949-y.

26. Dubrac S, Boneca IG, Poupel O, Msadek T. New insights into the WalK/WalR (YycG/YycF) essential signal transduction pathway reveal a major role in controlling cell wall metabolism and biofilm formation in S*taphylococcus aureus*. J Bacteriol. 2007;189(22):8257–69. doi: 10.1128/JB.00645-07.

27. Bleul L, Francois P, Wolz C. Two-Component Systems of *S. aureus*: Signaling and Sensing Mechanisms. Genes (Basel). 2021;13(1). doi: 10.3390/genes13010034.

28. Falord M, Mäder U, Hiron A, Debarbouille M, Msadek T. Investigation of the *Staphylococcus aureus* GraSR regulon reveals novel links to virulence, stress response and cell wall signal transduction pathways. PLoS One. 2011;6(7):e21323. doi: 10.1371/journal.pone.0021323.

29. Koprivnjak T, Mlakar V, Swanson L, Fournier B, Peschel A, Weiss JP. Cation-induced transcriptional regulation of the dlt operon of *Staphylococcus aureus*. J Bacteriol. 2006;188(10):3622–30. doi: 10.1128/JB.188.10.3622-3630.2006.

30. Seidl K, Leemann M, Zinkernagel AS. The ArlRS two-component system is a regulator of *Staphylococcus aureus*-induced endothelial cell damage. Eur J Clin Microbiol Infect Dis. 2018;37(2):289–92. doi: 10.1007/s10096-017-3130-5.

31. Kwiecinski JM, Crosby HA, Valotteau C, Hippensteel JA, Nayak MK, Chauhan AK, et al. *Staphylococcus aureus* adhesion in endovascular infections is controlled by the ArlRS-MgrA signaling cascade. PLoS Pathog. 2019;15(5):e1007800. doi: 10.1371/journal.ppat.1007800.

32. Walker JN, Crosby HA, Spaulding AR, Salgado-Pabon W, Malone CL, Rosenthal CB, et al. The *Staphylococcus aureus* ArlRS two-component system is a novel regulator of agglutination and pathogenesis. PLoS Pathog. 2013;9(12):e1003819. doi: 10.1371/journal.ppat.1003819.

33. Crosby HA, Schlievert PM, Merriman JA, King JM, Salgado-Pabon W, Horswill AR. The *Staphylococcus aureus* Global Regulator MgrA Modulates Clumping and Virulence by Controlling Surface Protein Expression. PLoS Pathog. 2016;12(5):e1005604. doi: 10.1371/journal.ppat.1005604.

34. Kwiecinski JM, Kratofil RM, Parlet CP, Surewaard BGJ, Kubes P, Horswill AR. Staphylococcus aureus uses the ArlRS and MgrA cascade to regulate immune evasion during skin infection. Cell Rep. 2021;36(4):109462. doi: 10.1016/j.celrep.2021.109462.

35. Radin JN, Kelliher JL, Parraga Solorzano PK, Kehl-Fie TE. The Two-Component System ArlRS and Alterations in Metabolism Enable *Staphylococcus aureus* to Resist Calprotectin-Induced Manganese Starvation. PLoS Pathog. 2016;12(11):e1006040. doi: 10.1371/journal.ppat.1006040.

36. de Baaij JH, Hoenderop JG, Bindels RJ. Magnesium in man: implications for health and disease. Physiol Rev. 2015;95(1):1–46. doi: 10.1152/physrev.00012.2014.

37. Fuchs S, Mehlan H, Bernhardt J, Hennig A, Michalik S, Surmann K, et al. AureoWiki ̵ The repository of the *Staphylococcus aureus* research and annotation community. Int J Med Microbiol. 2018;308(6):558–68. doi: 10.1016/j.ijmm.2017.11.011.

38. Fey PD, Endres JL, Yajjala VK, Widhelm TJ, Boissy RJ, Bose JL, et al. A genetic resource for rapid and comprehensive phenotype screening of nonessential *Staphylococcus aureus* genes. mBio. 2013;4(1):e00537–12. doi: 10.1128/mBio.00537-12.

39. Brown EJ, Flygare J, Hazenbos WL, Lehar SM, Mariathasan S, Morisaki JH, inventors; GENENTECH, INC, assignee. Anti-wall teichoic antibodies and conjugates.2014.

40. Pang YY, Schwartz J, Thoendel M, Ackermann LW, Horswill AR, Nauseef WM. agr-Dependent interactions of *Staphylococcus aureus* USA300 with human polymorphonuclear neutrophils. J Innate Immun. 2010;2(6):546–59. doi: 10.1159/000319855.

41. Crosby HA, Tiwari N, Kwiecinski JM, Xu Z, Dykstra A, Jenul C, et al. The *Staphylococcus aureus* ArlRS two-component system regulates virulence factor expression through MgrA. Mol Microbiol. 2020;113(1):103–22. doi: 10.1111/mmi.14404.

42. Sobhanifar S, Worrall LJ, Gruninger RJ, Wasney GA, Blaukopf M, Baumann L, et al. Structure and mechanism of *Staphylococcus aureus* TarM, the wall teichoic acid alpha-glycosyltransferase. Proc Natl Acad Sci U S A. 2015;112(6):E576–85. doi: 10.1073/pnas.1418084112.

43. Sobhanifar S, Worrall LJ, King DT, Wasney GA, Baumann L, Gale RT, et al. Structure and Mechanism of *Staphylococcus aureus* TarS, the Wall Teichoic Acid beta-glycosyltransferase Involved in Methicillin Resistance. PLoS Pathog. 2016;12(12):e1006067. doi: 10.1371/journal.ppat.1006067.

44. Burgui S, Gil C, Solano C, Lasa I, Valle J. A Systematic Evaluation of the Two-Component Systems Network Reveals That ArlRS Is a Key Regulator of Catheter Colonization by *Staphylococcus aureus*. Front Microbiol. 2018;9:342. doi: 10.3389/fmicb.2018.00342.

45. Luong TT, Dunman PM, Murphy E, Projan SJ, Lee CY. Transcription Profiling of the mgrA Regulon in *Staphylococcus aureus*. J Bacteriol. 2006;188(5):1899–910. doi: 10.1128/jb.188.5.1899-1910.2006.

46. Lei MG, Jorgenson MA, Robbs EJ, Black IM, Archer-Hartmann S, Shalygin S, et al. Characterization of Ssc, an N-acetylgalactosamine-containing *Staphylococcus aureus* surface polysaccharide. J Bacteriol. 2024;206(5):e0004824. doi: 10.1128/jb.00048-24.

47. Manna AC, Ingavale SS, Maloney M, van Wamel W, Cheung AL. Identification of sarV (SA2062), a new transcriptional regulator, is repressed by SarA and MgrA (SA0641) and involved in the regulation of autolysis in *Staphylococcus aureus*. J Bacteriol. 2004;186(16):5267–80. doi: 10.1128/JB.186.16.5267-5280.2004.

48. Jahnen-Dechent W, Ketteler M. Magnesium basics. Clin Kidney J. 2012;5(Suppl 1):i3-i14. doi: 10.1093/ndtplus/sfr163.

49. Ebel H, Günther T. Magnesium metabolism: a review. J Clin Chem Clin Biochem. 1980;18(5):257–70. doi: 10.1515/cclm.1980.18.5.257.

50. Urish KL, Cassat JE. *Staphylococcus aureus* Osteomyelitis: Bone, Bugs, and Surgery. Infect Immun. 2020;88(7). doi: 10.1128/IAI.00932-19.

51. Reffuveille F, Josse J, Velard F, Lamret F, Varin-Simon J, Dubus M, et al. Bone Environment Influences Irreversible Adhesion of a Methicillin-Susceptible *Staphylococcus aureus* Strain. Front Microbiol. 2018;9:2865. doi: 10.3389/fmicb.2018.02865.

52. Garcia-Betancur JC, Goni-Moreno A, Horger T, Schott M, Sharan M, Eikmeier J, et al. Cell differentiation defines acute and chronic infection cell types in *Staphylococcus aureus*. Elife. 2017;6:e28023. doi: 10.7554/eLife.28023.

53. Gryllos I, Grifantini R, Colaprico A, Jiang S, Deforce E, Hakansson A, et al. Mg(2+) signalling defines the group A streptococcal CsrRS (CovRS) regulon. Mol Microbiol. 2007;65(3):671–83. doi: 10.1111/j.1365-2958.2007.05818.x.

54. Hatoum-Aslan A. The phages of staphylococci: critical catalysts in health and disease. Trends Microbiol. 2021;29(12):1117–29. doi: 10.1016/j.tim.2021.04.008.

55. Moller AG, Lindsay JA, Read TD. Determinants of Phage Host Range in Staphylococcus Species. Appl Environ Microbiol. 2019;85(11):e00209–19. doi: 10.1128/AEM.00209-19.

56. Glowacka-Rutkowska A, Gozdek A, Empel J, Gawor J, Zuchniewicz K, Kozinska A, et al. The Ability of Lytic Staphylococcal Podovirus vB_SauP_phiAGO1.3 to Coexist in Equilibrium With Its Host Facilitates the Selection of Host Mutants of Attenuated Virulence but Does Not Preclude the Phage Antistaphylococcal Activity in a Nematode Infection Model. Front Microbiol. 2018;9:3227. doi: 10.3389/fmicb.2018.03227.

57. Monk IR, Shah IM, Xu M, Tan MW, Foster TJ. Transforming the untransformable: application of direct transformation to manipulate genetically *Staphylococcus aureus* and *Staphylococcus epidermidis*. mBio. 2012;3(2). doi: 10.1128/mBio.00277-11.

58. Weber BS, Ly PM, Feldman MF. Screening for Secretion of the Type VI Secretion System Protein Hcp by Enzyme-Linked Immunosorbent Assay and Colony Blot. Methods Mol Biol. 2017;1615:465–72. doi: 10.1007/978-1-4939-7033-9_32.

59. Hendriks A, van Dalen R, Ali S, Gerlach D, van der Marel GA, Fuchsberger FF, et al. Impact of Glycan Linkage to *Staphylococcus aureus* Wall Teichoic Acid on Langerin Recognition and Langerhans Cell Activation. ACS Infect Dis. 2021;7(3):624–35. doi: 10.1021/acsinfecdis.0c00822.

60. Prados J, Linder P, Redder P. TSS-EMOTE, a refined protocol for a more complete and less biased global mapping of transcription start sites in bacterial pathogens. BMC Genomics. 2016;17(1):849. doi: 10.1186/s12864-016-3211-3.

61. Goerke C, Koller J, Wolz C. Ciprofloxacin and trimethoprim cause phage induction and virulence modulation in *Staphylococcus aureus*. Antimicrob Agents Chemother. 2006;50(1):171–7. doi: 10.1128/AAC.50.1.171-177.2006.

62. Hanske J, Schulze J, Aretz J, McBride R, Loll B, Schmidt H, et al. Bacterial Polysaccharide Specificity of the Pattern Recognition Receptor Langerin Is Highly Species-dependent. J Biol Chem. 2017;292(3):862–71. doi: 10.1074/jbc.M116.751750.

63. Kropinski AM, Mazzocco A, Waddell TE, Lingohr E, Johnson RP. Enumeration of Bacteriophages by Double Agar Overlay Plaque Assay. In: Clokie MRJ, Kropinski AM, editors. Bacteriophages: Methods and Protocols, Volume 1: Isolation, Characterization, and Interactions. Totowa, NJ: Humana Press; 2009. p. 69–76.

64. Bowring JZ, Su Y, Alsaadi A, Svenningsen SL, Parkhill J, Ingmer H. Screening for Highly Transduced Genes in *Staphylococcus aureus* Revealed Both Lateral and Specialized Transduction. Microbiol Spectr. 2022;10(1):e0242321. doi: 10.1128/spectrum.02423-21.

65. Baba T, Takeuchi F, Kuroda M, Yuzawa H, Aoki K, Oguchi A, et al. Genome and virulence determinants of high virulence community-acquired MRSA. Lancet. 2002;359(9320):1819–27. doi: 10.1016/s0140-6736(02)08713-5.

66. Charpentier E, Anton AI, Barry P, Alfonso B, Fang Y, Novick RP. Novel cassette-based shuttle vector system for gram-positive bacteria. Appl Environ Microbiol. 2004;70(10):6076–85. doi: 10.1128/AEM.70.10.6076-6085.2004.

67. de Jong NW, van der Horst T, van Strijp JA, Nijland R. Fluorescent reporters for markerless genomic integration in *Staphylococcus aureus*. Sci Rep. 2017;7:43889. doi: 10.1038/srep43889.

68. Oduor JMO, Kiljunen S, Kadija E, Mureithi MW, Nyachieo A, Skurnik M. Genomic characterization of four novel Staphylococcus myoviruses. Arch Virol. 2019;164(8):2171–3. doi: 10.1007/s00705-019-04267-0.

69. Whittard E, Redfern J, Xia G, Millard A, Ragupathy R, Malic S, et al. Phenotypic and Genotypic Characterization of Novel Polyvalent Bacteriophages With Potent In Vitro Activity Against an International Collection of Genetically Diverse *Staphylococcus aureus*. Front Cell Infect Microbiol. 2021;11:698909. doi: 10.3389/fcimb.2021.698909.

